# Ribonucleotide synthesis by NME6 fuels mitochondrial gene expression

**DOI:** 10.1101/2022.11.29.518352

**Authors:** Nils Grotehans, Lynn McGarry, Hendrik Nolte, Moritz Kroker, Álvaro Jesús Narbona-Pérez, Soni Deshwal, Patrick Giavalisco, Thomas Langer, Thomas MacVicar

## Abstract

Replication and expression of the mitochondrial genome depend on the sufficient supply of nucleotide building blocks to mitochondria. Dysregulated nucleotide metabolism is detrimental to mitochondrial genomes and can result in instability of mitochondrial DNA and inflammation. Here, we report that a mitochondrial nucleoside diphosphate kinase, NME6, supplies mitochondria with ribonucleotides to drive the transcription of mitochondrial genes. Moreover, NME6 supports the maintenance of mitochondrial DNA when the access to cytosolic deoxyribonucleotides is limited. Perturbation of NME6 leads to the depletion of mitochondrial transcripts, destabilisation of the electron transport chain and impaired oxidative phosphorylation; deficiencies which are suppressed upon supplementation with pyrimidine ribonucleotides. Our work proposes NME6 and mitochondrial nucleotide metabolism to be untapped therapeutic targets in diseases associated with aberrant mitochondrial gene expression including cancer and autoimmune disorders.

## Introduction

Oxidative phosphorylation (OXPHOS) drives the synthesis of ATP during aerobic respiration and regulates broad cellular functions including redox homeostasis and cell death ^1^. The OXPHOS protein complexes at the inner mitochondrial membrane are predominantly composed of nuclear DNA encoded subunits that are imported into mitochondria. However, the correct assembly and activity of complexes I, III, IV and V also depend on the integration of subunits encoded by mitochondrial DNA (mtDNA) in the mitochondrial matrix ^2^. The compact and circular mtDNA encodes 13 OXPHOS subunits, 2 ribosomal RNAs (rRNAs) and 22 transfer RNAs (tRNAs) in mammalian cells and is maintained at a high, yet variable, copy number across tissues and developmental stages ^3^. Numerous processes such as mitochondrial dynamics and nucleotide metabolism regulate mtDNA replication and the abundance and quality of mtDNA. Defects in such mechanisms result in a broad spectrum of metabolic diseases ^4,5^.

The replication of mtDNA and synthesis of mitochondrial RNA (mtRNA) require a constant supply of deoxyribonucleotide triphosphates (dNTPs) and ribonucleotide triphosphates (rNTPs), respectively ^6,7^. Mammalian cells synthesise dNTPs and rNTPs in the cytosol from multiple carbon and nitrogen sources in a high energy demanding *de novo* synthesis pathway or, in a process termed nucleotide salvage, from pre-existing (deoxy)nucleosides via a series of phosphorylation reactions within the cytosol or mitochondria ^8^. Mitochondria cannot synthesise nucleotides *de novo* and therefore depend on the import of dNTPs and rNTPs across the inner mitochondrial membrane or the import of precursor (deoxy)nucleosides prior to phosphorylation by mitochondrial nucleotide salvage kinases ^9,10^ (Fig. 1A).

**Figure 1.**
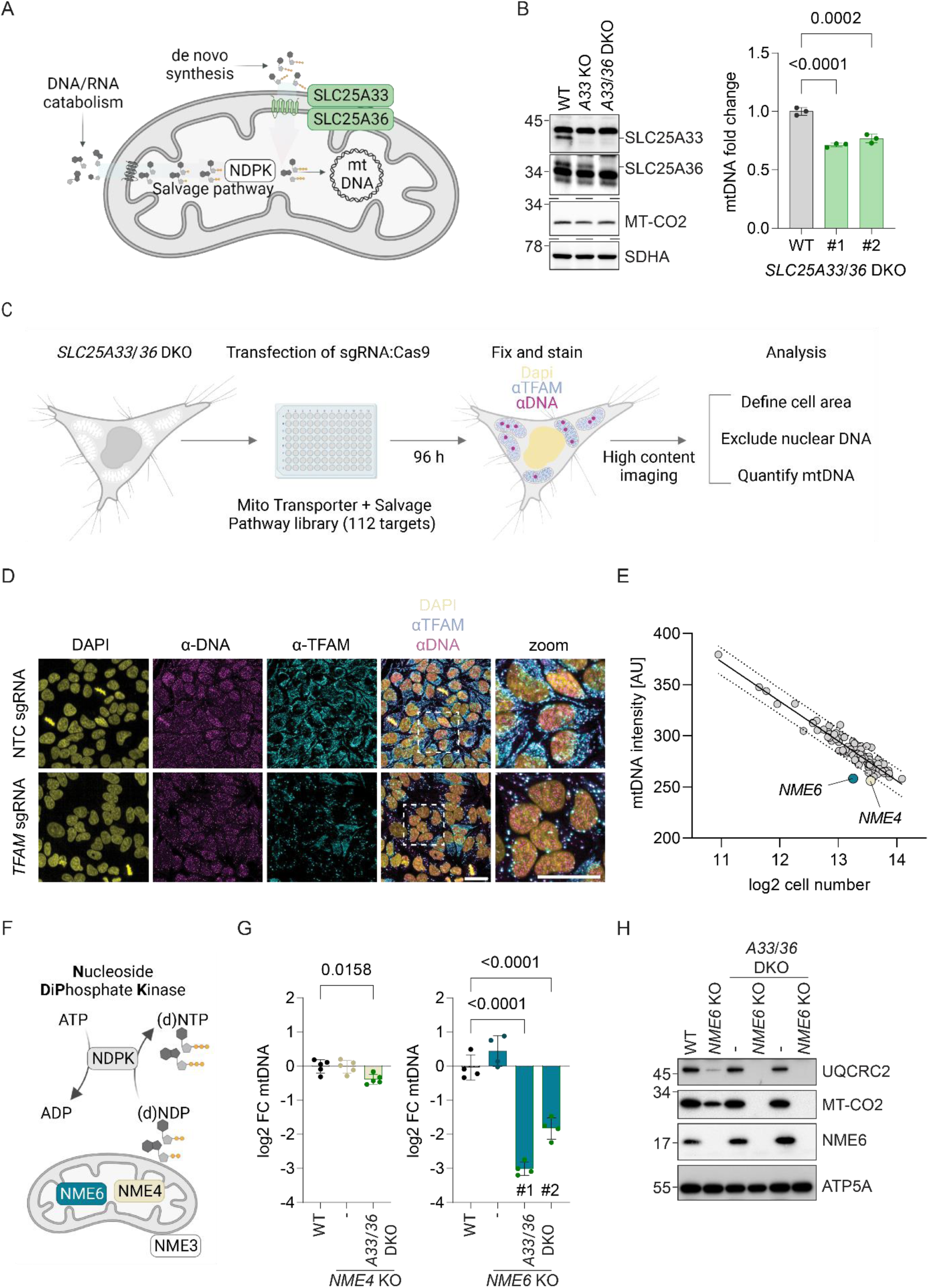
NME6 is required for the maintenance of mtDNA when mitochondrial pyrimidine import is blocked. (A) Scheme of the routes by which mitochondria obtain and metabolise pyrimidine nucleotides. (B) Immunoblot analysis of SLC25A33 and SLC25A36 depletion in the indicated knockout (KO) and double knockout (DKO) HeLa cells (left) alongside the relative mtDNA levels of two monoclonal *SLC25A33*/*SLC25A36* DKO cell lines calculated by qPCR (*CYTB / ACTB*; right) (n=3 independent cultures). (C) Experimental flow chart of the arrayed CRISPR-SpCas9 screen to identify regulators of mtDNA levels in the absence of mitochondrial pyrimidine import. (D) Representative images taken from the arrayed CRISPR-SpCas9 screen of *SLC25A33*/*SLC25A36* DKO cells showing one field of view from wells transfected with non-targeting control (NTC) sgRNA or TFAM sgRNA (scale bar = 20 µm) (E) The result of the CRISPR-SpCas9 screen plotted as the mean mtDNA intensity values against the mean log2 cell number from each sgRNA target. The values for cells transfected with *NME6* sgRNA and *NME4* sgRNA lie below the 95 % prediction bands (Least-squares regression; R^2^= 0.90; n=3 independent experiments; AU, arbitrary units). (F) Scheme of nucleoside diphosphate kinase (NDPK) enzymatic activity and the reported locations of the mitochondrial NDPKs: NME3, NME4 and NME6. (G) Relative mtDNA levels in *NME4* KO and *NME4*/*SLC25A33*/*SLC25A36* triple KO HeLa (left) or *NME6* KO and two clones of *NME6*/*SLC25A33*/*SLC25A36* triple KO HeLa (right). MtDNA calculated by qPCR (*CYTB / ACTB*) and presented as log2 fold change compared to levels in WT HeLa cells (n = 4-5 independent cultures). (H) Representative immunoblot analysis of the indicated HeLa cell lines (UQCRC2, Ubiquinol-cytochrome C reductase core protein 2; MT-CO2, mitochondrial encoded cytochrome c oxidase II; ATP5A, ATP synthase F1 subunit alpha). P values were calculated using one-way analysis of variance (ANOVA) with Tukey’s multiple comparison test (B, G). FC, fold change. Data are means ± standard deviation (SD).

The predominant source and supply route of dNTPs for mtDNA replication is often defined by the cell cycle and tissue type ^9^. The mitochondria within proliferating cells import *de novo* synthesised dNTPs from the cytosol, while quiescent cells have a greater dependence on mitochondrial nucleotide salvage as a consequence of downregulated cytosolic dNTP synthesis ^10,11^. This is reflected, for instance, by the severe depletion of mtDNA in the skeletal muscle of patients with mutations in the mitochondrial pyrimidine salvage pathway enzyme thymidine kinase 2 (*TK2*) ^12,13^. However, this remains a sweeping generalisation and we understand very little surrounding the regulation of dNTP supply for mtDNA replication in fluctuating metabolic conditions, such as those encountered during the progression and treatment of cancer.

Recent work has highlighted that mitochondrial nucleotide metabolism impacts cellular nucleotide balance with striking consequences for cellular signalling. Enhanced uptake of mitochondrial pyrimidines can cause nuclear genomic instability due to a depletion of cytosolic dNTPs ^14^. On the other hand, it promotes mtDNA replication but can also trigger mtDNA release from mitochondria and mtDNA-dependent inflammatory pathways ^15^. The dNTP salvage pathway also contributes to innate immune signalling whereby upregulation of the mitochondrial cytidine/uridine monophosphate kinase 2 (CMPK2) drives the rapid synthesis of mtDNA in macrophages subjected to pro-inflammatory stimuli to support inflammasome activation ^16,17^.

The work introduced above provides evidence that adjusting mitochondrial dNTP supply can tune mtDNA replication, but we know very little regarding the regulation of rNTP supply for mitochondrial transcription. The mitochondrial RNA polymerase (POLRMT) incorporates rNTPs to transcribe the polycistronic RNA in addition to non-coding 7S RNA, which is an abundant mtRNA transcript that functions in a feedback loop to negatively regulate POLRMT activity and RNA primers required for mtDNA replication ^18^. Mitochondrial transcription is therefore recognised as a crucial process in OXPHOS regulation. Upregulation of mitochondrial transcription machinery, including POLRMT and the transcription elongation factor (TEFM) have been reported in several cancers including non-small cell lung cancer and hepatocellular carcinoma ^19–21^ and mitochondrial transcription is an exciting potential therapeutic target in cancer ^22^.

Here, we demonstrate that regulated mitochondrial ribonucleotide metabolism is critical for mitochondrial gene expression. By interrogating mitochondrial nucleotide supply pathways, we reveal that the poorly characterised nucleoside diphosphate kinase, NME6, is a mitochondrial nucleotide salvage pathway enzyme that is essential for the maintenance of mtDNA when mitochondrial pyrimidine nucleotide import is blocked. Strikingly, NME6 is required for constitutive mtRNA synthesis and OXPHOS function in proliferating cells, which highlights a crucial role for mitochondrial ribonucleotide supply in mitochondrial transcription. We propose that targeting nucleotide supply via NME6 represents a new way to modulate mitochondrial transcription.

## Results

To address how nucleotide supply regulates mitochondrial gene maintenance and expression, we first explored the routes by which mitochondria obtain cytosolic pyrimidine nucleotides for mtDNA and mtRNA synthesis in proliferating cells. We decided to focus on the mitochondrial supply of pyrimidines since we and others have observed that enhanced mitochondrial import or salvage of pyrimidines is sufficient to increase the abundance of mtDNA ^15,16,23^. While several mitochondrial solute carriers are known to exchange adenine nucleotides across the inner membrane, only two mitochondrial pyrimidine (deoxy)nucleotide carriers (SLC25A33 and SLC25A36) have been identified in mammalian cells ^24,25^, both of which also transport guanine nucleotides *in vitro* ^24^. Defective mitochondrial uptake of guanine nucleotides was recently associated with hyperinsulinism/hyperammonemia syndrome in patients with deleterious mutation of *SLC25A36* ^26^. Loss of the single homologue of *SLC25A33* and *SLC25A36* in yeast, Rim2, leads to mtDNA depletion and blocks growth on non-fermentable carbon sources ^27^. We generated HeLa cells lacking *SLC25A33* and *SLC25A36* by CRISPR-SpCas9 mediated genome editing and monitored mtDNA levels. To our surprise, mtDNA levels were only reduced by 20 % in these cells (Fig. 1B), suggesting to us that mitochondria obtain pyrimidines via an alternative route when the pyrimidine nucleotide carriers are missing.

### The maintenance of mtDNA depends on NME6 when pyrimidine import is blocked

We hypothesised that alternative mitochondrial carriers and/or the mitochondrial nucleotide salvage pathway can supply pyrimidines in the absence of SLC25A33 and SLC25A36. To test this, we performed an arrayed CRISPR-SpCas9 knockout high-content microscopy screen to identify genes that regulate mtDNA content in cells lacking the canonical mitochondrial pyrimidine transporters. HeLa cells lacking both pyrimidine nucleotide carriers were transfected with SpCas9 nuclease and an arrayed CRISPR library wherein each well contained three sgRNAs targeting individual genes with proposed roles in mitochondrial metabolite transport or pyrimidine nucleotide salvage (112 genes) (Table S1). The mtDNA was visualised 96 h following transfection by immunofluorescence with an anti-DNA antibody and we quantified the mean fluorescence intensity per cell after exclusion of nuclear DNA signal (Fig. 1C, D). The mtDNA level inversely correlated with cell confluency in each well, which led us to plot the mean fluorescence intensity of mtDNA against cell number to identify potential outliers (Fig. 1E). Two outliers corresponded to sgRNA targeting the mitochondrial nucleoside diphosphate kinases (NDPK), *NME4* and *NME6*, which suggested that their loss renders cells unable to maintain mtDNA levels when pyrimidine import is blocked (Fig. 1E). The library also included sgRNA targeting mitochondrial transcription factor A (TFAM), a high-mobility group (HMG)- box domain containing transcription factor ^28^ and mtDNA-binding protein that is essential for the packaging of mtDNA into compact nucleoids ^29,30^. TFAM protein level is known to correlate closely with the abundance of mtDNA, for example *Tfam* heterozygous knockout mouse tissues lose approximately 50 % mtDNA while TFAM overexpression increases mtDNA levels ^31–34^. We were therefore surprised that *TFAM* sgRNA did not register as an outlier in our mtDNA fluorescence intensity analysis (Fig 1E). Selective depletion of TFAM in cells transfected with *TFAM* sgRNA was confirmed in the screen using an anti-TFAM antibody (Fig. 1D). The residual TFAM condensed with mtDNA in enlarged nucleoids in line with previous observations in cells transiently depleted of TFAM ^35,36^. As this might explain why TFAM was not detected as an outlier in our initial analysis, we also calculated the area of mtDNA puncta as an additional readout of mtDNA homeostasis. We found that, analogous to the mean fluorescence intensity, the total area of mtDNA also declined with cell confluency and cells transfected with *TFAM* sgRNA emerged with strongly reduced mtDNA area per cell as expected. A modest reduction in mtDNA area was also observed in cells transfected with *NME6* sgRNA (Fig. S1A).

The results from our CRISPR screen indicated that NME4 and NME6 may maintain mtDNA in the absence of pyrimidine nucleotide transport. The non-metastatic (NME) gene family of NDPKs generate (d)NTPs by transferring the terminal phosphate group predominantly from ATP to dNDPs or rNDPs via a transient phospho-histidine intermediate (Fig. 1F) ^37^. Three NME family members are reported to reside at mitochondria in different subcompartments; NME3 is located at the mitochondrial surface ^38^, NME4 has been reported to be exposed to both the intermembrane space and matrix ^39,40^ and NME6 is imported to the mitochondrial matrix ^41^. Since the reduction of mtDNA upon transfection with *NME4* and *NME6* sgRNA was subtle (Fig 1E), we generated stable monoclonal knockout cells by plasmid expression of *NME4* or *NME6* sgRNA and determined mtDNA levels by real-time quantitative PCR (qPCR). NME4 loss did not strongly alter mtDNA levels in the presence or absence of SLC25A33 and SLC25A36 (Fig. 1G), which suggests that NME4 is not required for pyrimidine nucleotide salvage in these cells. Conversely, while NME6 depletion alone did not reduce mtDNA levels, the combined knockout of *NME6*, *SLC25A33* and *SLC25A36* caused a dramatic loss of mtDNA to 15-25 % of WT levels (Fig. 1G) and resulted in the loss of the mtDNA-encoded protein cytochrome c oxidase II (MT-CO2; Fig. 1H). To further validate our findings, we depleted SLC25A33 and SLC25A36 individually and together in WT and *NME6* knockout cells using short interfering RNA (esiRNA). Knockdown of the pyrimidine carriers had little effect on mtDNA levels in WT cells but significantly depleted mtDNA in cells lacking NME6 (Fig. S1B, C). NME6 is ubiquitously expressed in humans and catalyses phosphotransfer through a conserved histidine residue within an NDPK consensus motif at position 137 (H137) ^42^ and kinase independent functions of NME6 have also been proposed ^41,43^. Importantly, mtDNA levels were maintained in *NME6* knockout cells expressing WT NME6-MycFlag but not in cells expressing kinase inactive mutant NME6 (NME6^H137N^-MycFlag) (Fig. S1B). Collectively, these data demonstrate that NME6 maintains the mitochondrial genome upon reduced pyrimidine nucleotide supply from the cytosol and indicate that NME6 catalyses the final enzymatic step in the salvage of nucleotides within mitochondria (Fig 1A).

### NME6 supports cell proliferation independent of mtDNA synthesis

Our finding that mtDNA levels are unchanged in cells depleted of NME6 alone implied that NME6 is redundant in mitochondria that obtain sufficient dNTPs from the cytosol. However, analysis of DepMap CRISPR knockout screens highlighted a fitness dependency of many cancer cell lines on *NME6* (Fig. 2A) and we found that the loss of NME6 limits the growth of HeLa cells in normal glucose medium that was restored upon the expression of NME6-MycFlag (Fig. 2B). The growth defect was more severe in galactose medium, which forces HeLa cells to depend on glutaminolysis and oxidative phosphorylation (OXPHOS) for energy supply ^44,45^ (Fig. 2C). *NME6* was recently identified in a CRISPR-SpCas9 screen as essential for survival in human plasma like medium (HPLM) ^46^ and, consistently, we observed that NME6 depleted HeLa cells could not survive in HPLM for extended periods (Fig. 2D). This was likely due to glucose exhaustion in the HPLM of *NME6* knockout HeLa cells since glucose supplementation could restore the viability of NME6-depleted cells in HPLM (Fig S2A). These growth assays revealed that NME6 supports cell proliferation, particularly in respiratory-dependent conditions. The increased dependency on NME6 in galactose or HPLM medium was independent of mtDNA levels, which remained normal in cells lacking NME6 regardless of the growth medium (Fig. 2E, F). Our results thus demonstrate that proliferating cells require NME6, even when the mitochondrial import of dNTPs is normal and the maintenance of mtDNA does not depend on NME6.

**Figure 2.**
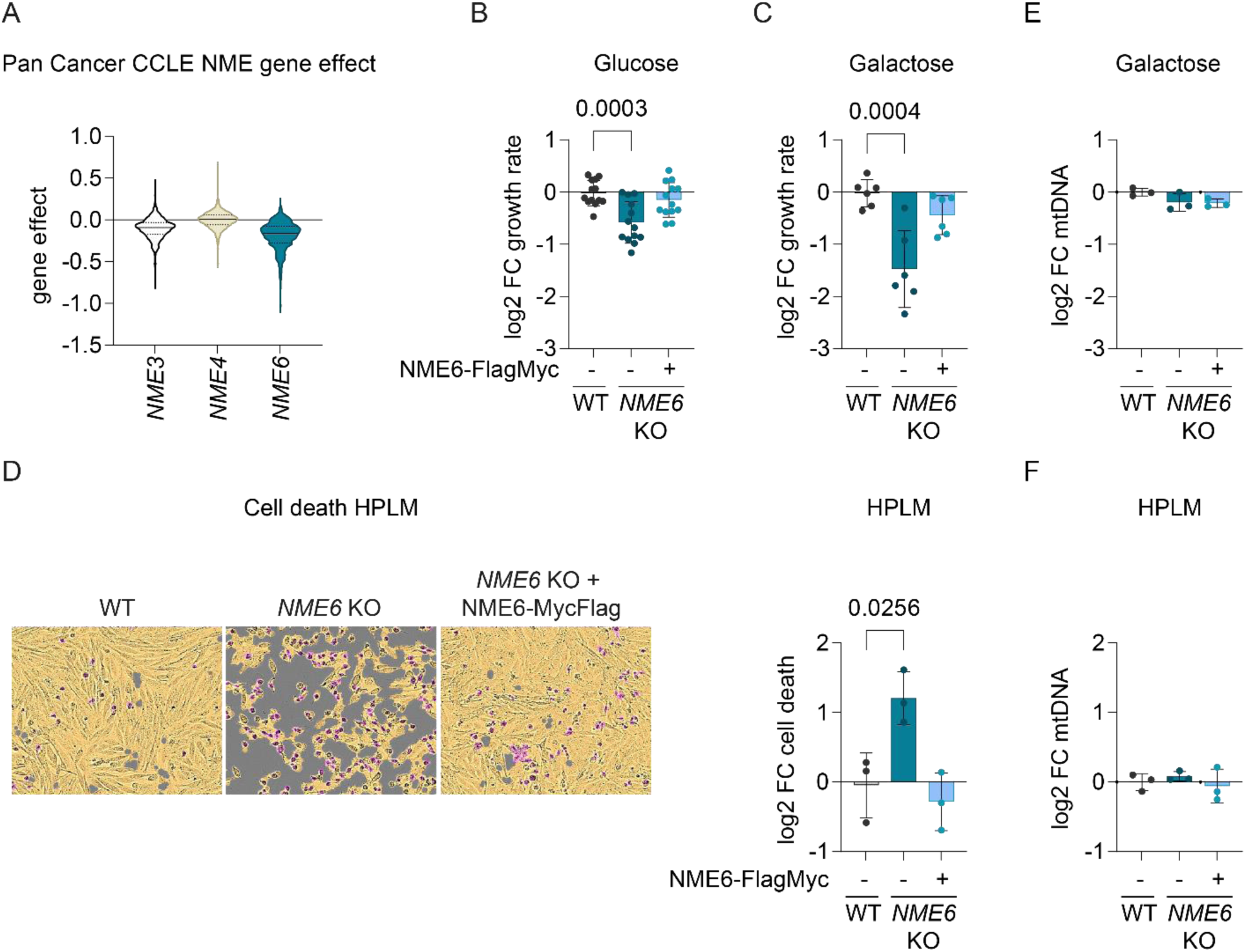
NME6 supports cell proliferation independent of mtDNA synthesis. (A) Violin plot of gene effects of *NME3*, *NME4* or *NME6* depletion in 1086 cell lines from the Cancer Cell Line Encyclopaedia (CCLE) determined by CRISPR screening (DepMap 22Q2 Public+Score, Chronos; solid line denotes median, dotted line denotes 25 % quartile). (B) The growth rate of *NME6* KO and *NME6* KO + NME6-MycFlag HeLa relative to WT HeLa cells incubated in DMEM containing 25 mM glucose (log2; n=13 independent cultures). (C) The growth rate of *NME6* KO and *NME6* KO + NME6-MycFlag HeLa relative to WT HeLa cells incubated in DMEM containing 10 mM galactose (log2; n=6 independent cultures). (D) Representative live-cell images of the indicated cell lines grown in Human Plasma Like Medium (HPLM) (left) and calculated cell death after 96 h (right). Cell confluency is depicted with the yellow mask and dead cells are identified by SYTOX green staining in purple (n=3 independent cultures). (E) MtDNA level monitored by qPCR (*CYTB / ACTB*) in *NME6* KO and *NME6* KO + NME6-MycFlag HeLa relative to WT HeLa cells in DMEM containing 10 mM galactose (n=3 independent cultures). (F) MtDNA level monitored by qPCR (*CYTB / ACTB*) in *NME6* KO and *NME6* KO + NME6-MycFlag HeLa relative to WT HeLa cells in HPLM (n=3 independent cultures). P values were calculated using one-way ANOVA with Tukey’s multiple comparison test (B, C, D, E, F). FC, fold change. Data (except A) are means ± SD.

### Mitochondrial respiration and OXPHOS subunit homeostasis depend on NME6

We sought to explain why NME6 is required for cell proliferation and therefore investigated its influence on mitochondrial function. Consistent with a growth defect in galactose medium (Fig. 2C), we found that the loss of NME6 strongly reduced cellular oxygen consumption rates (OCR) and resulted in a concomitant increase in extracellular acidification rate (ECAR) (Fig. 3A). Enhanced ECAR is indicative of upregulated glycolysis, which likely explains the greater glucose dependency in cells lacking NME6 (Fig S2A). Normal OCR and ECAR were restored in *NME6* knockout cells upon the expression of WT NME6-MycFlag but not NME6^H137N^-MycFlag, demonstrating that mitochondrial function depends on NME6 kinase activity (Fig. 3A).

**Figure 3.**
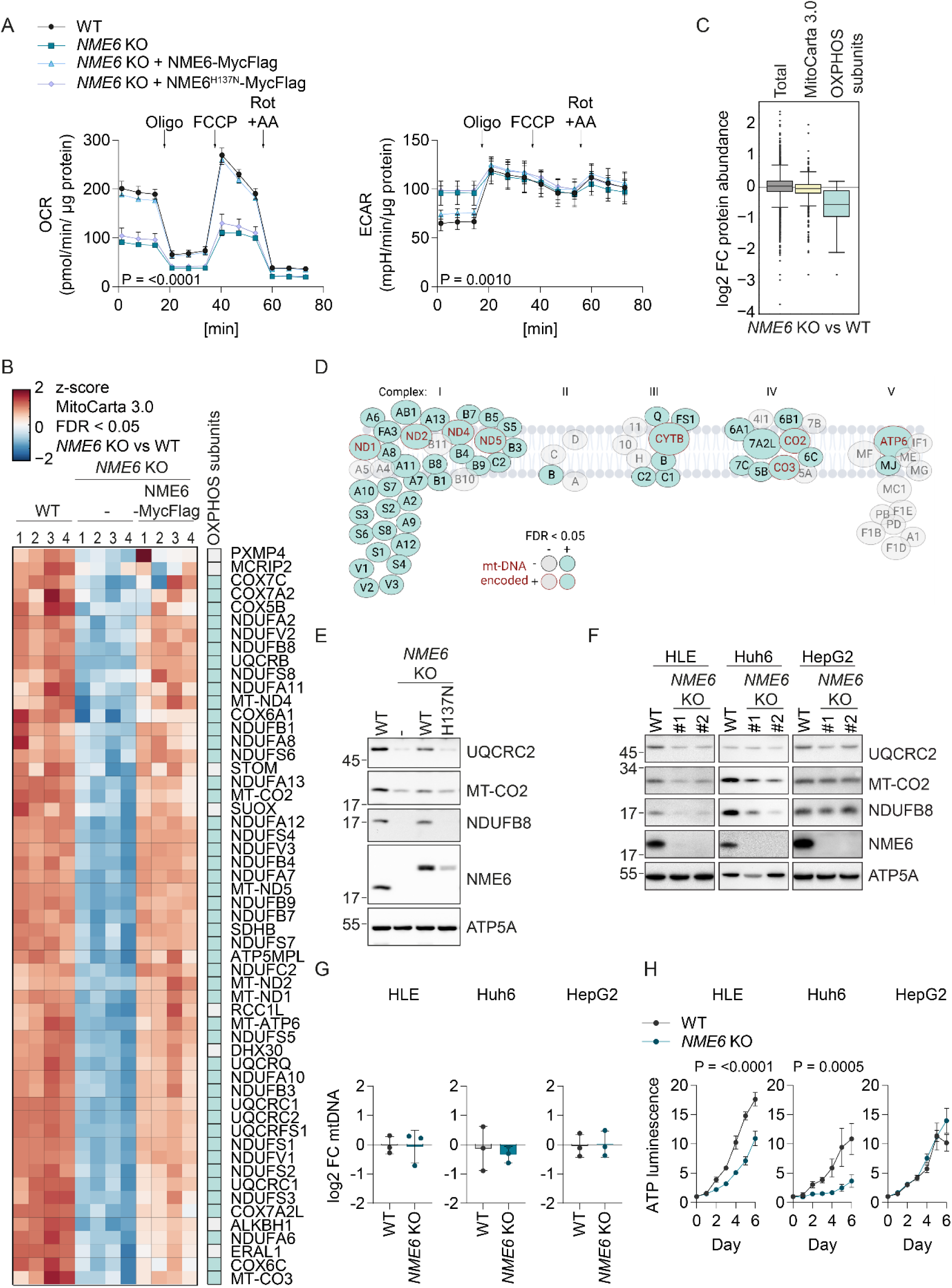
Mitochondrial respiration and OXPHOS depend on NME6. (A) Oxygen consumption rates (OCR) and extracellular acidification rates (ECAR) of the indicated HeLa cell lines during a mitochondrial stress test with inhibitor treatments at the indicated timepoints (Oligo, oligomycin; FCCP, carbonyl cyanide-p-trifluoromethoxyphenylhydrazone; Rot+AA, rotenone and antimycin a; n=3 independent experiments). (B) Heat map representation of z-scores of log2-transformed protein intensities determined by quantitative mass spectrometry and filtered for mitochondrial proteins according to MitoCarta 3.0 ^47^ (cluster five of Fig. S3A). The proteins shown clustered together due to their significant (permutation-based FDR < 0.05, s_0_ = 0.1) depletion in *NME6* KO HeLa cells compared to WT and *NME6* KO + NME6- MycFlag HeLa cells. OXPHOS subunits are indicated in the right column (n=4 independent cultures). (C) Box plot analysis of log2 fold change in protein intensities in *NME6* KO compared to WT HeLa cells. Distribution of the complete set of quantified protein groups (Total), the MitoCarta 3.0 positive as well as the set of OXPHOS subunits (CI – CV of the MitoCarta 3.0 pathway annotations. (Box limits denote 25- and 75 % quartile, line denotes the median, whiskers denote 1.5 x interquartile range deviation from the median). (D) Graphical representation of all respiratory complex subunit proteins detected in our proteomic assay to highlight the respiratory complexes most affected by loss of NME6 as also shown in B. OXPHOS subunits depleted in *NME6* KO cells compared to WT and *NME6* KO + NME6-MycFlag HeLa cells are in teal. MtDNA-encoded OXPHOS subunits are labelled in red. Subunits that were not significantly altered between genotypes are in grey. (E) Immunoblot analysis of WT HeLa cells, *NME6* KO cells and *NME6* KO cells expressing NME6-MycFlag (WT) or NME6^H137N^-MycFlag (H137N). (F) Immunoblot analysis of WT cells and two *NME6* KO clones (#1, #2) generated in three different liver cancer cell lines: HLE, Huh6 and HepG2. (G) MtDNA level monitored by qPCR (*CYTB / ACTB*) in *NME6* KO relative WT cells in the indicated liver cancer cell lines (n=3 independent cultures). (H) The relative growth of WT and *NME6* KO liver cancer cell lines monitored on each day (d) using an ATP luminescence assay (n=4 independent cultures). P values were calculated using two-way ANOVA (A, H) and unpaired t-test (G). P value for two-way ANOVA factor: genotype is indicated. Multiple testing correction by permutation-based false discovery rate (FDR) estimated to 0.05; FC, fold change. Data (except C) are means ± SD.

To explore why mitochondrial respiration depends on NME6, we examined mitochondrial protein homeostasis by quantitative proteomics and performed unsupervised hierarchical clustering analysis of mitochondrial proteins (MitoCarta 3.0)^47^ that were significantly altered in an NME6-dependent manner (Fig. S3A). The largest cluster consisted of 55 mitochondrial proteins that were significantly depleted in cells lacking NME6 compared to WT and NME6-MycFlag complemented cells (Fig. 3B). Remarkably, 47 of the 55 mitochondrial proteins were OXPHOS subunits, resulting in the collective depletion of OXPHOS proteins relative to other mitochondrial proteins, which was also revealed by unbiased 1D-enrichment driven pathway analysis (Fig. 3C). These data demonstrate that OXPHOS subunit homeostasis depends on NME6, while overall mitochondrial mass is unaffected by NME6 loss. Specifically, the majority of complex I, III and IV subunits were reduced in the absence of NME6 and included all the mtDNA-encoded OXPHOS subunits measured (Fig. 3D). Immunoblotting of selected OXPHOS subunits revealed that OXPHOS protein homeostasis could not be restored in *NME6* knockout cells complemented with the kinase dead NME6^H137N^-MycFlag (Fig. 3E), which is consistent with the respiratory dependency on phospho-active NME6 in these cells (Fig. 3A). While OXPHOS proteins were decreased, we observed the accumulation of >20 proteins in *NME6* knockout cells (Fig. S3A, B). These proteins included known targets of the integrated stress response (ISR) and components of the one-carbon metabolism and of other metabolic pathways, which may be induced in response to the OXPHOS deficiency in cells lacking NME6^48,49^.

We next expanded our analysis to test the role of NME6 in liver cancer cell lines. We chose to study these cells since liver cancer cell lines display a significant increase in NME6 levels relative to other cancer types (Fig. S3C) and the expression of NME6 correlates with an unfavourable prognosis in liver cancer patients (Fig. S3D). OXPHOS subunits were diminished in two out of three NME6 knockout human liver cancer cell lines (Fig. 3F), despite no alteration in the level of mtDNA across all conditions (Fig. 3G). The absence of NME6 in HLE and Huh6 cells, but not HepG2 cells, also resulted in a significant growth defect (Fig. 3H), which was consistent with publicly available CRISPR screening data (Fig. S3E; depmap.org/portal) and correlated with the levels of OXPHOS subunits in these cell lines (Fig. 3F). Finally, complementation of *NME6* knockout HLE cells with WT NME6-MycFlag but not NME6^H137N^-MycFlag restored OXPHOS homeostasis (Fig. S3F), thus confirming the requirement for NME6 kinase activity in these cells. Together, we conclude that NME6 is required for respiration and the maintenance of OXPHOS subunits in proliferating cells.

### Mitochondrial gene expression depends on NME6

We wondered how a mitochondrial NDPK regulates the abundance of OXPHOS subunits, independent from its role in supporting mtDNA synthesis. We first searched for clues by identifying genes with similar effects to NME6 on cellular fitness ^50^. Gene coessentiality network analysis across hundreds of heterogenous cancer cell lines using the FIREWORKS (Fitness Interaction Ranked nEtWORKS) web tool ^51^ revealed that the top coessential genetic interactors with *NME6* are genes encoding mitochondrial proteins, which was unique within the NME gene family (Fig. 4A). Specifically, *NME6* dependency correlates with regulators of mtDNA replication, transcription, mitochondrial tRNA maturation and mitochondrial ribosome (mitoribosome) biogenesis (Fig. 4A). This analysis strongly indicated that NME6 is involved in the expression of mtDNA-encoded genes. We therefore monitored the synthesis rate of mtDNA-encoded proteins by determining the incorporation of ^35^S-methionine during *in vitro* mitochondrial translation. We observed reduced synthesis of the majority of mtDNA-encoded subunits in mitochondria lacking NME6 (Fig. 4B, C), which pointed to defective mitochondrial transcription or translation in the absence of NME6.

**Figure 4.**
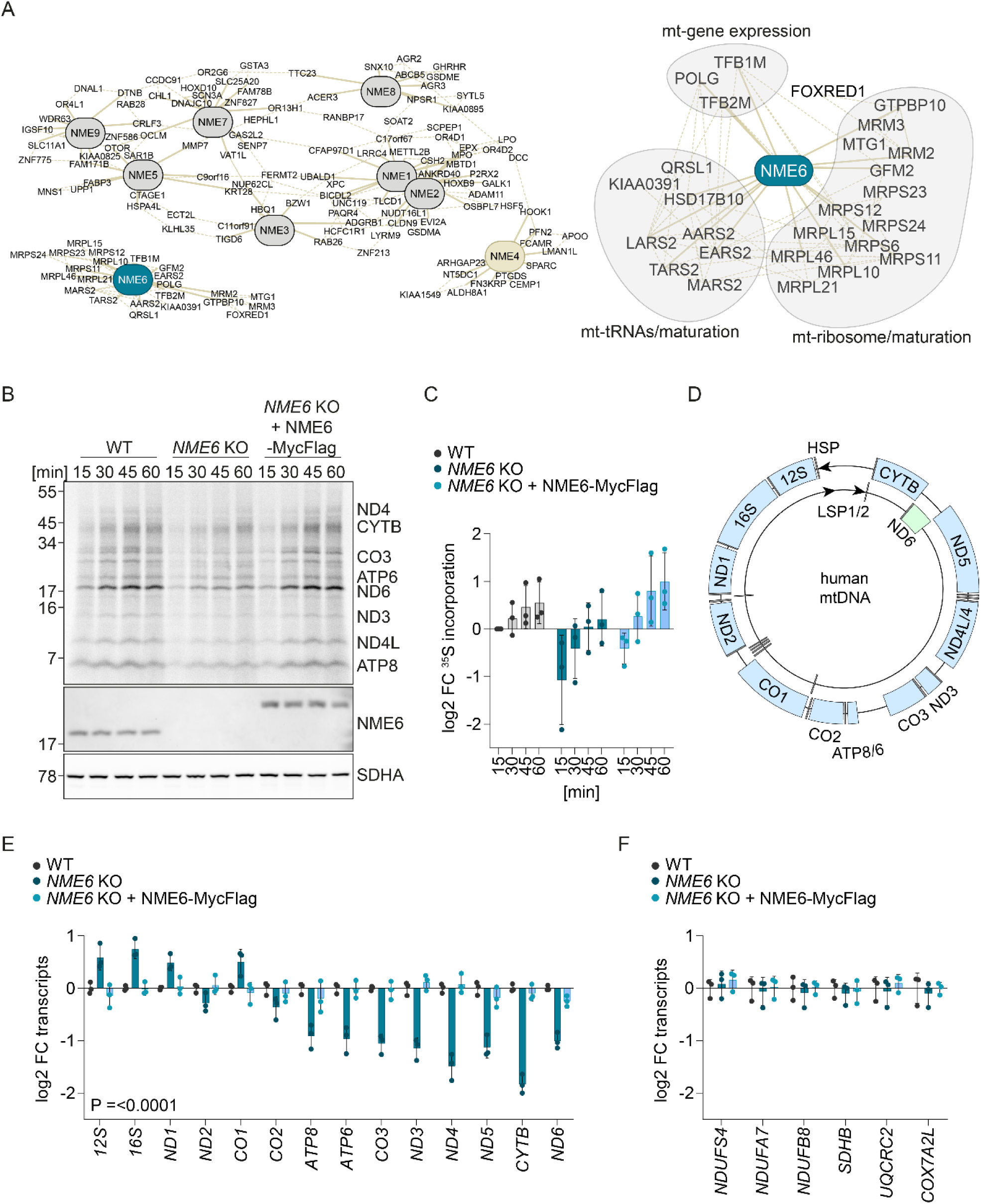
NME6 regulates mitochondrial gene expression. (A) Pan-cancer gene coessentiality network visualisation of the top 15 positively correlated genes (solid lines) and 5 secondary node positive interactions (dotted lines) with all NME family members (*NME1-9*) using FIREWORKS ^51^ (left) and *NME6* correlated genes were grouped further into manually annotated mitochondrial functions (right). (B) Representative mitochondrial translation assay monitored by the incorporation of ^35^S methionine and cysteine into the indicated mtDNA-encoded proteins followed by autoradiography (top panel). MtDNA encoded proteins are labelled according to expected size and NME6 and SDHA immunoblots are shown below. (C) Quantification of ^35^S methionine and cysteine incorporation into all mtDNA-encoded proteins labelled in (B) relative to WT HeLa cells at 15 min (log2; mitochondrial preparations from n=3 independent cultures). (D) Scheme of human mtDNA organisation with the heavy strand promoter (HSP) represented by an arrow on the outside and the light strand promoters (LSP) by two arrows on the inside. Heavy strand encoded subunits are labelled in blue, light strand encoded subunit is labelled in green, tRNAs are labelled in grey. (E) MtDNA-encoded transcript levels analysed by qRT-PCR in *NME6* KO and *NME6* KO + NME6- MycFlag cells relative to WT HeLa cells (log2; n=3 independent cultures). (F) Nuclear DNA-encoded transcript levels analysed by qRT-PCR in *NME6* KO and *NME6* KO + NME6- MycFlag HeLa cells relative to WT (log2; n=3 independent cultures). P values were calculated using two-way ANOVA (C, E, F). P value for two-way ANOVA factor: genotype is indicated FC, fold change. Data are means ± SD.

Immunoprecipitation of NME6-MycFlag coupled with mass spectrometry identified the putative mitochondrial ribosome assembly factor, RCC1L, to be the only high confidence interaction partner of NME6 (Fig. S4A), in line with previous proximity labelling and immunoprecipitation assays ^41,52,53^. However, unlike cells depleted of RCC1L ^54^, we did not observe any disruption of mitoribosome proteins in cells lacking NME6 (Fig. 3B, S3A), which argues against NME6 being required for mitoribosome assembly. Mitochondrial transcription is initiated from a single promoter on the heavy-strand and two promoters on the light-strands of mtDNA to yield polycistronic transcripts that are further processed to individual mitochondrial messenger (m)RNAs, transfer (t)RNAs and ribosomal (r)RNAs ^55,56^ (Fig. 4D). We measured the levels of mitochondrial mRNA and rRNA by real-time quantitative reverse transcription PCR (qRT-PCR) and observed a striking pattern of mitochondrial mRNA depletion in HeLa cells lacking NME6 that correlated with the distance from the heavy strand promoter (Fig. 4D, E). Heavy-strand RNA transcripts up to *ATP8* were barely affected and the levels of the promoter proximal *12S* and *16S* rRNA were higher in NME6-depleted cells (Fig. 4E). However, heavy-strand mRNAs from *ATP8* onwards, as well as *ND6* on the light-strand, were significantly lower in the absence of NME6 compared to WT and NME6-MycFlag complemented cells (Fig. 4E). Conversely, the transcript levels of nuclear DNA encoded OXPHOS subunits were unaffected by the loss of NME6 (Fig. 4F) despite their reduced protein levels (Fig. 3D). Collectively, these data highlight a crucial role for NME6 in the maintenance of mitochondrial RNA transcripts, which could explain the OXPHOS deficiency in cells lacking NME6.

### NME6 supplies rNTPs for mitochondrial transcription

To synthesise the almost genome-length polycistronic transcripts, POLRMT requires an adequate supply of rNTPs. NDPKs phosphorylate dNDPs or rNDPs with little specificity between nucleotide bases ^57^ and we therefore hypothesised that NME6 supplies rNTPs for mitochondrial transcription. To test this hypothesis, we supplemented *NME6* knockout cells with rNTPs or dNTPs and measured mitochondrial transcript levels by qPCR. Strikingly, mitochondrial mRNAs at lower steady state levels in *NME6* knockout cells were restored to WT levels upon treatment with rNTPs, while supplementation with dNTPs had no effect (Fig. 5A). Proteomic analysis confirmed that supplementation with rNTPs, was sufficient to increase the levels of OXPHOS subunits in *NME6* knockout cells (Fig. 5B). Exogenous rNTPs are likely hydrolysed by ectonucleotidases prior to cell uptake as ribonucleosides via concentrative and equilibrative nucleoside transporters ^58,59^. We therefore treated cells with a mix of precursor nucleosides that included the four ribonucleosides cytidine, uridine, guanosine and adenosine and the deoxyribonucleoside thymidine. Similar to rNTP treatment, nucleoside supplementation resulted in the complete rescue of mitochondrial transcript levels in cells lacking NME6 (Fig. 5C). We reasoned that imported nucleosides must be phosphorylated via the cytosolic salvage pathway prior to entry into the mitochondria in order to bypass the requirement for NME6. Indeed, we found that nucleoside treatment no longer restored mitochondrial transcript levels in *NME6* knockout cells that were depleted of the cytosolic uridine-cytidine kinase 2 (UCK2), which is essential for pyrimidine salvage by phosphorylating uridine and cytidine (Fig. 5D, S5A, B). Importantly, the rescue of mitochondrial transcripts by nucleoside treatment correlated with a complete restoration of normal OCR and ECAR in *NME6* knockout cells (Fig. 5E) which further demonstrated that exogenous nucleoside supply is sufficient to maintain OXPHOS in the absence of NME6.

**Figure 5.**
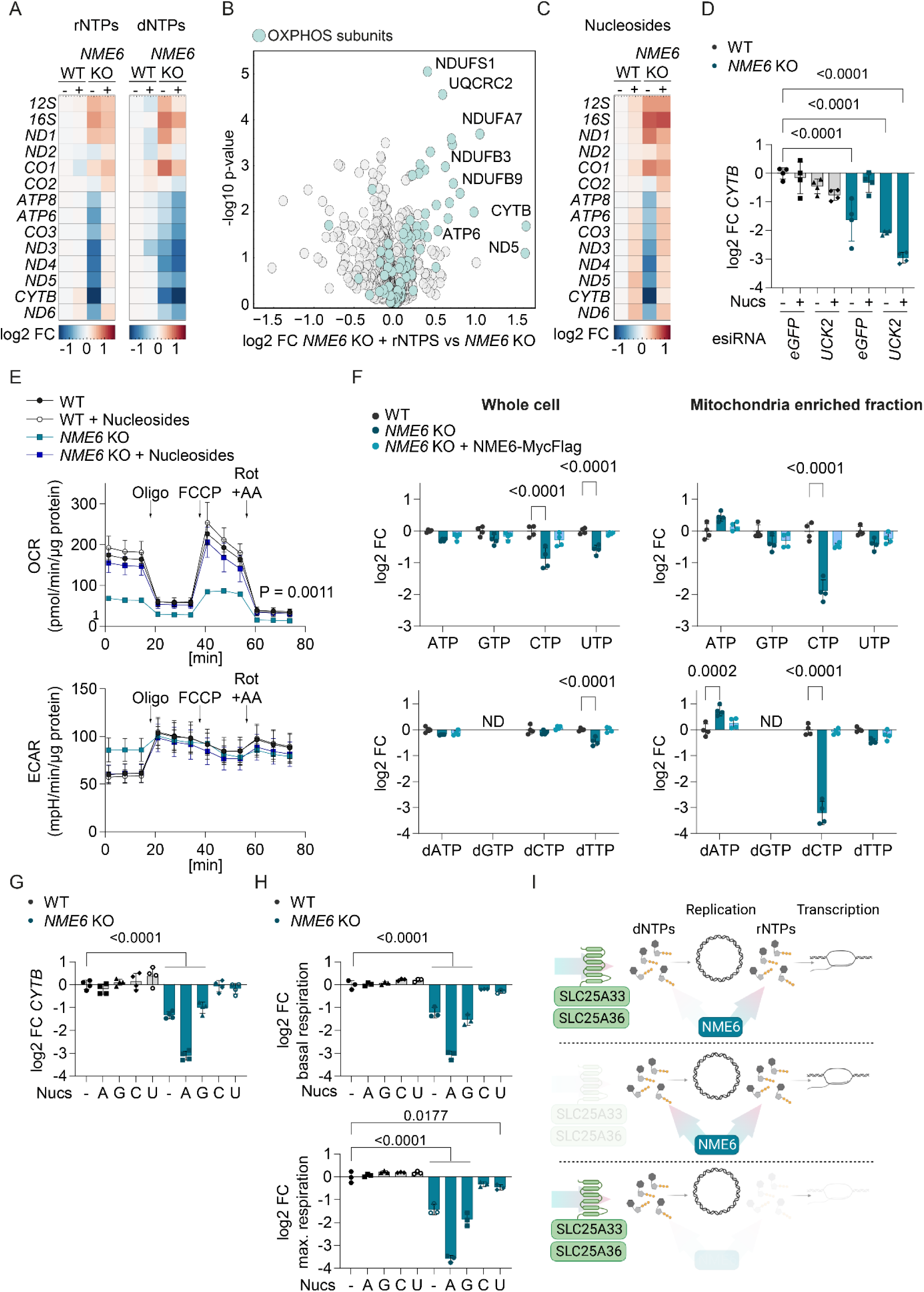
NME6 maintains pyrimidine ribonucleotide triphosphates for mitochondrial transcription. (A) Heat map of log2 transformed mean mitochondrial transcript levels of WT and *NME6* KO HeLa cells incubated with 100 µM rNTPs or dNTPs for 48 h relative to untreated WT cells analysed by qRT-PCR (n=3 independent cultures). (B) Volcano plot representation of log2 fold change in proteins between *NME6* KO cells treated with 100 µM rNTPs for 120 h and untreated *NME6* KO HeLa cells determined by quantitative mass spectrometry. OXPHOS subunits are highlighted in teal (n=3 independent cultures) (C) Heat map of log2 transformed mean mitochondrial transcript levels of WT and *NME6* KO HeLa cells incubated with 100 µM nucleosides for 48 h relative to untreated WT cells analysed by qRT-PCR (n=3 independent cultures). (D) *CYTB* transcript levels analysed by qRT-PCR in WT and *NME6* KO HeLa cells transfected with the indicated esiRNA and incubated with or without nucleosides (100 µM) for 72 h (log2; n=4 independent cultures). (E) Oxygen consumption rates (OCR) and extracellular acidification rates (ECAR) of WT and *NME6* KO HeLa cells incubated with or without nucleosides (100 µM) for a minimum of 120 h. Mitochondrial stress test was performed as in Fig. 3A (n=3 independent experiments). (F) Nucleotide levels in whole cell (left) and mitochondria enriched fractions (right) from *NME6* KO and *NME6* KO + NME6-MycFlag (WT) cells compared to WT HeLa cells as determined by quantitative mass spectrometry. The NTPs (top) and dNTPs (bottom) are shown except for dGTP which was not detected in our analysis. P values are shown where significant (log2; n=4 independent cultures). (G) *CYTB* transcript levels analysed by qRT-PCR in WT and *NME6* KO HeLa cells incubated with the indicated nucleoside species for 48 h (A, adenosine; G, guanosine; C, cytidine; U, uridine; log2; 100 µM; n=4 independent cultures) (H) Basal (top) and maximal (bottom) oxygen consumption rates of WT and *NME6* KO HeLa cells incubated with the indicated nucleoside species for a minimum of 120 h relative to untreated WT cells (A, adenosine; G, guanosine; C, cytidine; U, uridine; log2; 100 µM; n=4 independent experiments). Basal and maximal rates were calculated from Fig S5E. (I) NME6 is required to supply ribonucleotides (NTPs) for transcription and deoxynucleotides (dNTPs) for mtDNA replication if mitochondrial pyrimidine nucleotide uptake is blocked. Loss of NME6 therefore primarily leads to defective mitochondrial gene expression that can be accompanied by severe mtDNA depletion in the absence of mitochondrial pyrimidine import. P values were calculated using two-way ANOVA (E) with Tukey’s multiple comparison test (D, F) or one-way ANOVA with Tukey’s multiple comparison test (G, H). P value for two-way ANOVA factor: genotype is indicated. FC, fold change. Data are means ± SD.

Our nucleoside supplementation experiments strongly indicated that NME6 is required to generate rNTPs for mitochondrial transcription and OXPHOS. To strengthen this conclusion, we quantified the abundance of nucleotide species in whole-cell extracts and mitochondrial fractions by liquid chromatography-mass spectrometry (LC-MS; Fig. 5F, S5C). Mitochondria lacking NME6 had a dramatic depletion of CTP and dCTP and a mild reduction of UTP, which were restored in cells re-expressing NME6. Conversely, GTP levels were unchanged, while ATP and dATP were even moderately increased in mitochondria lacking NME6 (Fig. 5F). This result suggested that NME6 is predominantly required for the maintenance of mitochondrial pyrimidine triphosphates, some of which were also reduced in whole cell extracts (Fig. 5F). This likely explains why the knockdown of *UCK2* alone was sufficient to block the nucleoside rescue of mitochondrial transcripts in *NME6* knockout cells (Fig. 5D). These data compelled us to compare the effect of individual nucleoside species on mitochondrial transcript levels in NME6 knockout cells. Treatment of cells with cytidine or uridine restored *CYTB* transcripts in the absence of NME6, whereas purine nucleosides did not (Fig. 5G). Uridine treatment can increase both cellular UTP and CTP levels ^60^ since UTP is readily converted to CTP by the cytosolic enzyme CTP synthetase (CTPS). Finally, we tested the impact of individual nucleoside supplementation on mitochondrial respiration and observed that cytidine or uridine treatment resulted in near complete restoration of OCR in *NME6* knockout HeLa cells (Fig. 5H, S5E). The ECAR in *NME6* knockout cells was also normalised upon treatment with the pyrimidine nucleosides, likely reflecting a deceleration of glycolysis (Fig. 5H, S5E). Conversely, treatment of these cells with guanosine did not restore mitochondrial respiration or ECAR, while adenosine treatment resulted in a more severe inhibition of mitochondrial respiration and a further increase in ECAR in cells lacking NME6. The bioenergetic impact of each nucleoside correlated remarkably with their individual effects on mitochondrial transcript levels in *NME6* knockout cells (Fig. 5G, S5D) and nucleoside treatment had no impact on the bioenergetics (Fig. 5H S5E) or mitochondrial transcript levels in WT HeLa cells (Fig 5G, S5D). These results highlight a specific dependency on NME6 for the maintenance of mitochondrial pyrimidine nucleotides required to drive mitochondrial gene expression and OXPHOS in proliferating cells.

## Discussion

We demonstrate an unexpected regulatory role of the mitochondrial nucleotide diphosphate kinase NME6 for mitochondrial transcription. As part of the mitochondrial nucleotide salvage pathway, NME6 provides rNTPs to POLRMT, which limits mitochondrial transcription and OXPHOS activity in proliferating cells. Moreover, NME6 is required for the maintenance of mtDNA and provides dNTPs for mtDNA replication when pyrimidine import from the cytosol is limited. Our results thus reveal two independent roles of NME6 for the maintenance and expression of the mitochondrial genome.

The loss of NME6 kinase activity results in severe OXPHOS deficiency due to the reduced expression of mtDNA-encoded genes and subsequent disruption of OXPHOS subunits, which likely reflects the degradation of non-assembled OXPHOS subunits upon a primary loss of mtDNA encoded subunits ^61,62^. Our results are also consistent with previous reports that complex V subunits remain relatively stable in comparison to complex I, III and IV subunits when mtDNA gene expression is compromised ^63,64^. The depletion of mtDNA-encoded transcripts in the absence of NME6 occurs despite normal levels of mtDNA and mitochondrial nucleotide transporters that allow the uptake of cytosolic nucleotides. The majority of mitochondrial transcripts are depleted in cells lacking NME6, apart from the rRNAs and the four mRNAs most proximal to the HSP (ND1, ND2, CO1 and CO2). The restoration of mitochondrial transcript levels in NME6 depleted cells by treatment with exogenous nucleosides or ribonucleotides is strong evidence that NME6 is a primary supplier of rNTPs for mitochondrial transcription. The depletion of rNTPs may lead to the stalling of the mitochondrial RNA polymerase, analogous to DNA polymerase stalling upon the depletion of dNTPs, and ultimately perturb transcription efficiency and incompletion of polycistronic transcripts ^65^. The different effects of NME6 loss on mitochondrial transcripts likely reflects progressive rNTP depletion with ongoing transcription and argues against a general role for NME6 in mRNA stabilisation. For instance, all heavy-strand mRNA transcripts are depleted in mitochondria lacking the mRNA stabilising factor, leucine-rich pentatricopeptide repeat containing (LRPPRC) protein ^66,67^. Interestingly, our proteomic analysis revealed that the levels of four mitochondrial proteins associated with mitochondrial RNA granules (MRGs) are reduced in the absence of NME6: ALKBH1, ERAL1, DHX30 and RCC1L ^68–71^. MRGs are hubs for the processing and maturation of nascent RNA and are associated with the assembly of mitoribosomes ^72^. We confirmed that NME6 interacts with RCC1L but we did not observe any disruption of mitoribosome proteins in cells lacking NME6, which argues against NME6 being required for mitoribosome assembly ^54^. Nevertheless, in light of the critical role of NME6 for transcription, it is intriguing to consider that its complex formation with RCC1L may allow spatial coordination of mitochondrial transcription with translation.

Our metabolomics data demonstrated that the accumulation of mitochondrial pyrimidines, particularly CTP and dCTP, is dependent on NME6 in HeLa cells. This was surprising given the reported lack of base moiety specificity of NDPK enzymes ^57^. Interestingly, *in vitro* kinase assays that describe NME6 as an active NDPK used CDP as the γ-phosphate acceptor with recombinant NME6 ^42^, whereas no NDPK activity was detected for recombinant NME6 when dTDP was used as the γ-phosphate acceptor ^41^. While the specific enzymatic activity of NME6 remains to be dissected and is out of the scope of this study, it is also possible that alternative pathways support the maintenance of purine rNTPs in the absence of NME6. Indeed, the fact that NME6 was dispensable for mitochondrial gene expression and proliferation in HepG2 cells points to the likely existence of alternative metabolic routes that may compensate NME6 deficient cells with mitochondrial rNTPs.

Our CRISPR-SpCas9 screen revealed that *NME6* is also required for the maintenance of mtDNA in cells lacking the mitochondrial pyrimidine transporters, SLC25A33 and SLC25A36. It is likely that mtDNA replication depends on dNTP production by NME6 when mitochondrial pyrimidine import is blocked. Alternatively, the loss of mtDNA may result from severe transcription deficiency in mitochondria lacking pyrimidine transporters and NME6 since mtDNA replication requires the synthesis of RNA primers by POLRMT ^73,74^. The screen also indicated that NME4 regulates mtDNA levels in cells lacking mitochondrial pyrimidines. However, further experiments did not reveal any effect on mtDNA levels in NME4-depleted cells regardless of mitochondrial pyrimidine import. NME4 was the first mitochondrial NDPK to be characterised ^40^ and has been proposed to support the synthesis of mtDNA ^16,75–78^ in addition to other functions that include supplying the mitochondrial fusion GTPase, OPA1, with GTP in the intermembrane space ^79,80^. While evidence to support a direct role for NME4 in the maintenance of steady-state mtDNA levels is lacking, NME4 was reported to support the stimulation of mtDNA replication and the cytosolic release of oxidised mtDNA in macrophages, leading to activation of the inflammasome ^16^. Similarly, NME6 was recently identified as a positive regulator of the inflammasome in a mouse macrophage cell line, along with NME4 and NME3 ^17^. It is possible that the contribution of mitochondrial NDPKs to mtDNA synthesis differs depending on cell and tissue type and that compensatory activity may exist between mitochondrial NDPKs, which may explain why mutation of NDPKs are not associated with mitochondrial DNA depletion syndromes. It remains to be seen whether NME6 is essential for mtDNA synthesis in quiescent tissues that suppress *de novo* synthesis of nucleotides and depend on mitochondrial nucleotide salvage for the provision of dNTPs for mtDNA synthesis ^10,11,81^.

In summary, we demonstrate that NME6 is required for mitochondrial transcription and OXPHOS activity. Tuning mitochondrial nucleotide supply via NME6 thus represents a new approach to modulate mitochondrial gene expression. Expanding our understanding of mitochondrial nucleotide metabolism further will be essential to understand how mitochondrial nucleotide supply impacts ageing and disease associated with dysregulation of the mitochondrial genome and OXPHOS function. NME6 is also a promising novel target for the treatment of certain cancers that are expected to respond to the inhibition of POLRMT and mitochondrial transcription ^22^.

## Acknowledgements

We thank the MPI Biology of Ageing Metabolomics core facility, the Beatson Advanced Imaging Resource and D. Diehl for technical assistance. We thank Dusanka Milenkovic, Payam Gammage, Vanessa Xavier and Ryan Corbyn for helpful discussion and Catherine Winchester for comments on the manuscript. We thank Victor Villar and Saverio Tardito for providing liver cancer cell lines. This work was supported by funds of the Max-Planck-Society and from the Deutsche Forschungsgemeinschaft (DFG, German Research Foundation) - SFB1403 – Projektnummer 414786233) to T.L. and a Cancer Research UK Career Development Fellowship to TM (RCCFELCDF-May21\100001) and CRUK core funding to the CRUK Beatson Institute (A31287).

## Supplementary Information

**Supplementary Figures S1-S5**

Table S1 (related to Figure 1). Data and sgRNA sequences from CRISPR-SpCas9 screen

Table S2 (related to Figure 3). *NME6* KO HeLa proteomics

Table S3 (related to Figure S4). NME6-FLAG immunoprecipitation proteomics

Table S4 (related to Figure 5). *NME6* KO HeLa proteomics +/- rNTPs

### Supplementary Figures

**Figure S1.**
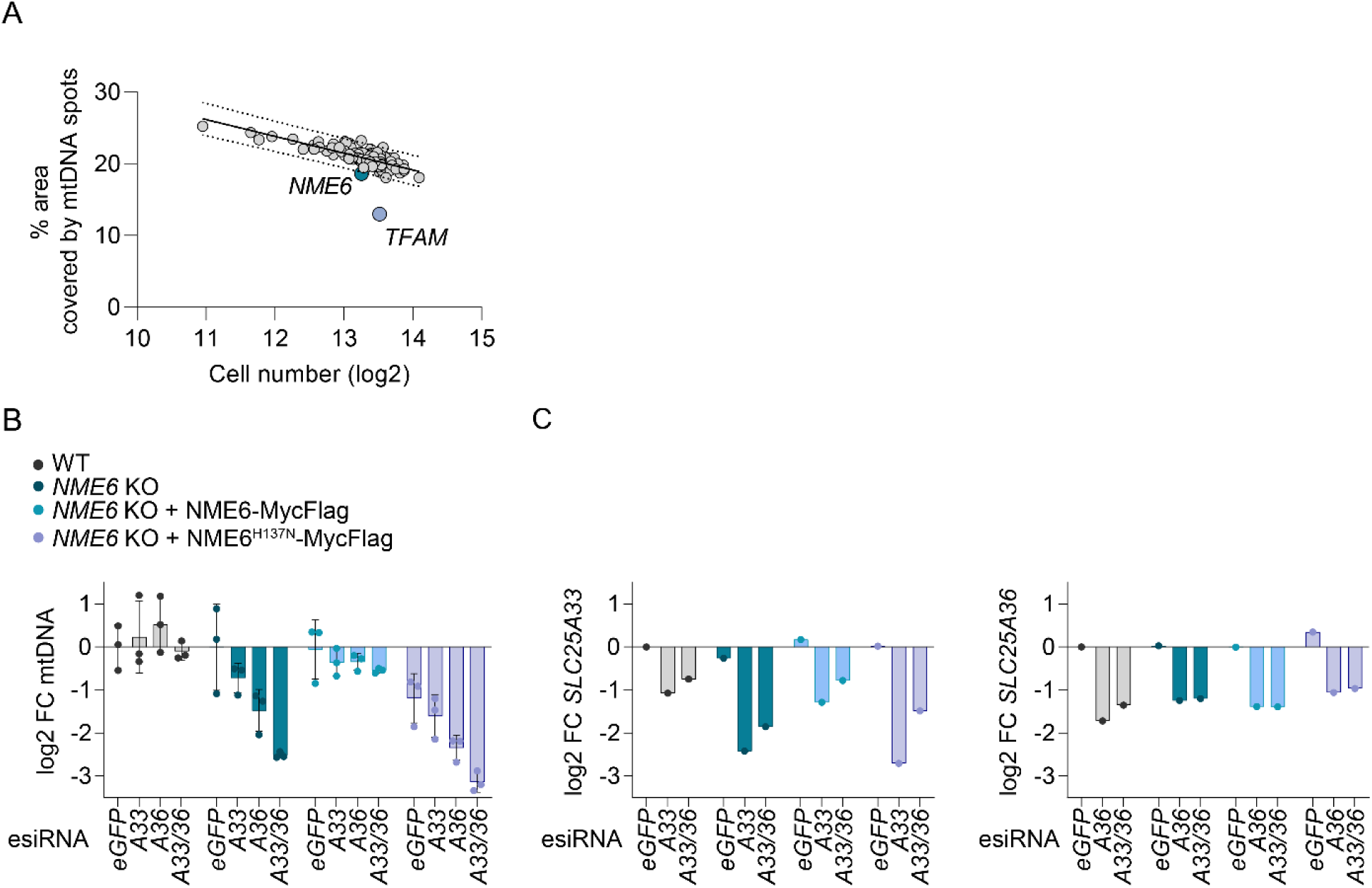
(related to Figure 1). CRISPR-SpCas9 screen analysis and validation by RNA interference. (A) The result of the CRISPR-SpCas9 screen (Fig. 1C) plotted as the percentage of cell cytoplasm area occupied by mtDNA spots against the mean log2 cell number from each sgRNA target. The values for cells transfected with *NME6* sgRNA and *TFAM* sgRNA lie below the 95 % prediction bands (Least-squares regression; R^2^= 0.52; n=3 independent experiments). (B) MtDNA level monitored by qPCR (*CYTB / ACTB*) in the indicated HeLa cell lines treated with esiRNA targeting *GFP* (control), *SLC25A33* (A33), *SLC25A36* (A36) or *SLC25A33* + *SLC25A36* (A33/A36) relative to WT cells (log2; n=3 independent cultures). (C) *SLC25A33* (left) and *SLC25A36* (right) transcript levels in samples from a representative experiment in B monitored by qRT-PCR (log2; n=1) FC, fold change. Data are means ± SD.

**Figure S2.**
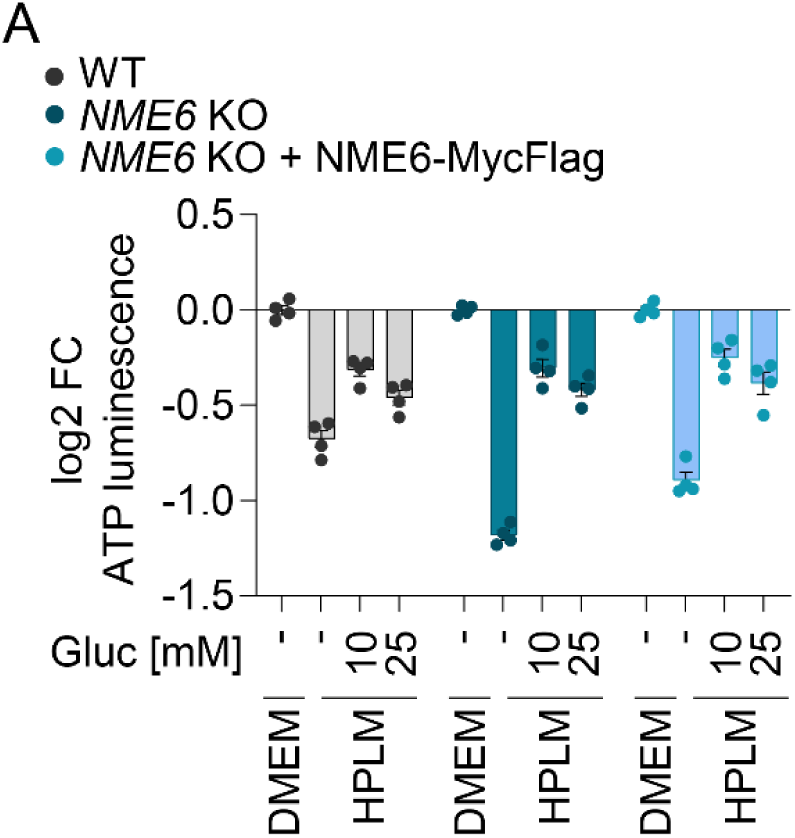
(related to Figure 2). Glucose is limiting for growth in HPLM. (A) Cell viability in WT, *NME6* KO and *NME6* KO + NME6 MycFlag HeLa cells incubated in HPLM supplemented with different concentrations of glucose (standard HPLM contains 5 mM glucose). Cell viability was determined by ATP luminescence assay and analysed relative to DMEM (log 2; n=4 independent cultures). FC, fold change. Data are means ± SD.

**Figure S3.**
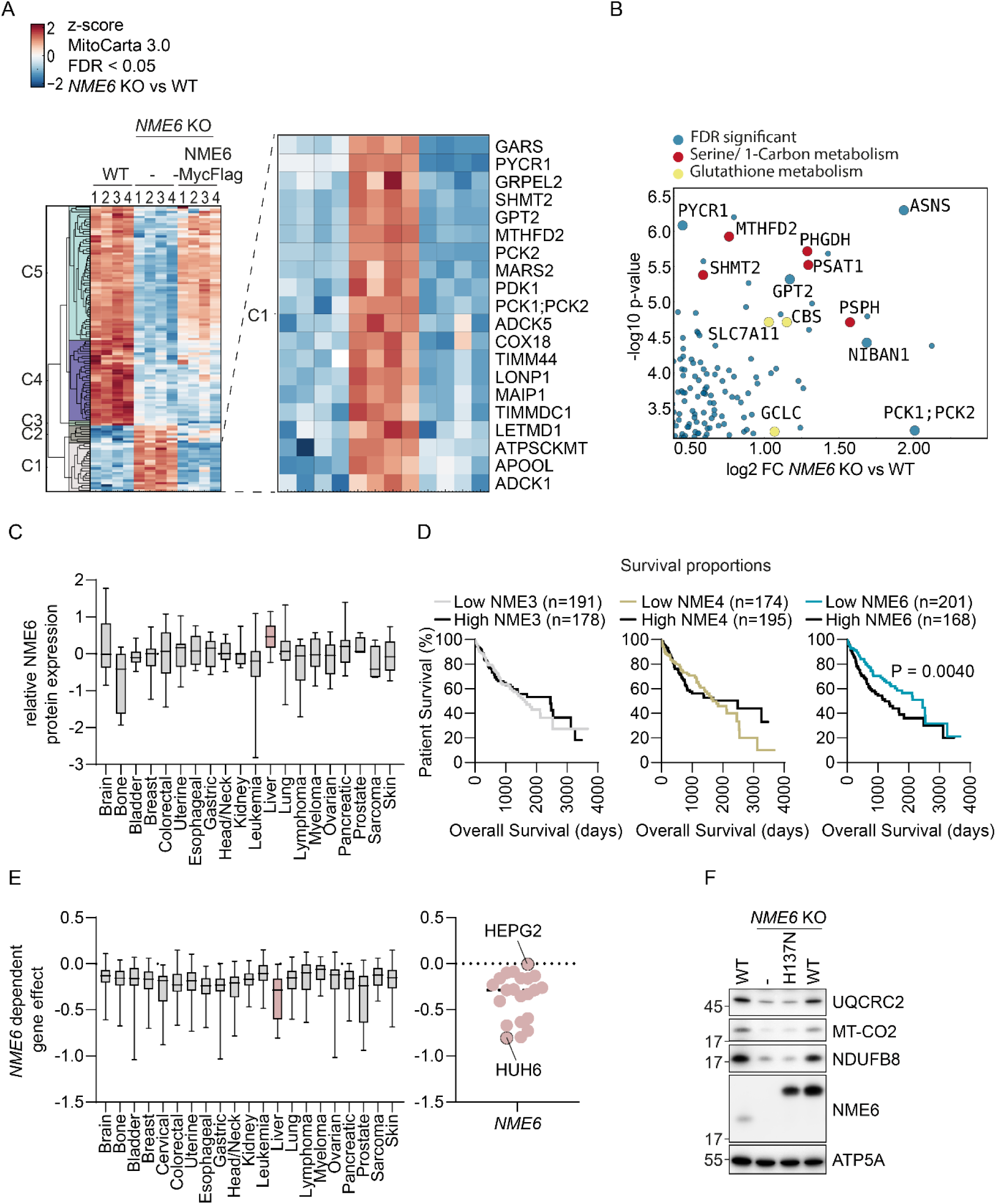
(related to Figure 3). Alterations to the proteome upon NME6 depletion and further characterisation of NME6 in liver cancer. (A) Extended unsupervised hierarchical row clustering (Euclidean distance, complete method) representation of significantly different poritens (FDR < 0.05) (*NME6* KO vs WT) z-scores of log2-transformed protein intensities determined by quantitative mass spectrometry and filtered for mitochondrial proteins according to MitoCarta 3.0 ^47^. The cluster presented in Fig. 3B is visible at the top (C5). The bottom cluster (C1) is expanded here to reveal proteins that are significantly upregulated in *NME6* KO cells compared to WT and *NME6* KO + NME6-MycFlag HeLa cells (n=4 independent cultures). (B) Volcano plot representation of all proteins that significantly accumulate in *NME6* KO relative to WT HeLa cells from the same experiment as in A and Fig. 3B, C. Proteins involved in serine / 1 carbon metabolism and glutathione metabolism are indicated (n=4 independent cultures). (C) Relative NME6 protein expression in cell lines across the indicated cancer types determined by quantitative proteomic profiling ^82^. Data obtained from depmap.org/portal (solid line = median, box limits = 25^th^ and 75^th^ percentile and whiskers = maxima and minima). (D) Kaplan-Meier plots showing patient survival in the indicated liver hepatocellular carcinoma cohorts (LIHC) from The Cancer Genome Atlas Program (TCGA) analysed using http://www.tcga-survival.com. P values were determined using a log rank test ^83^. (E) Stratification of the gene effect of NME6 depletion from Fig. 2A into cancer type (left, solid line = median, box limits = 25^th^ and 75^th^ percentile and whiskers = maxima and minima). Individual cell line *NME6* gene effects are also shown with HepG2 and Huh6 cell lines highlighted (right, note that HLE cells are not included in DepMap 22Q2 Public+Score, Chronos) (F) Immunoblot analysis of WT HLE cells, *NME6* KO cells and *NME6* KO cells expressing NME6-MycFlag (WT) or NME6^H137N^-MycFlag (H137N).

**Figure S4.**
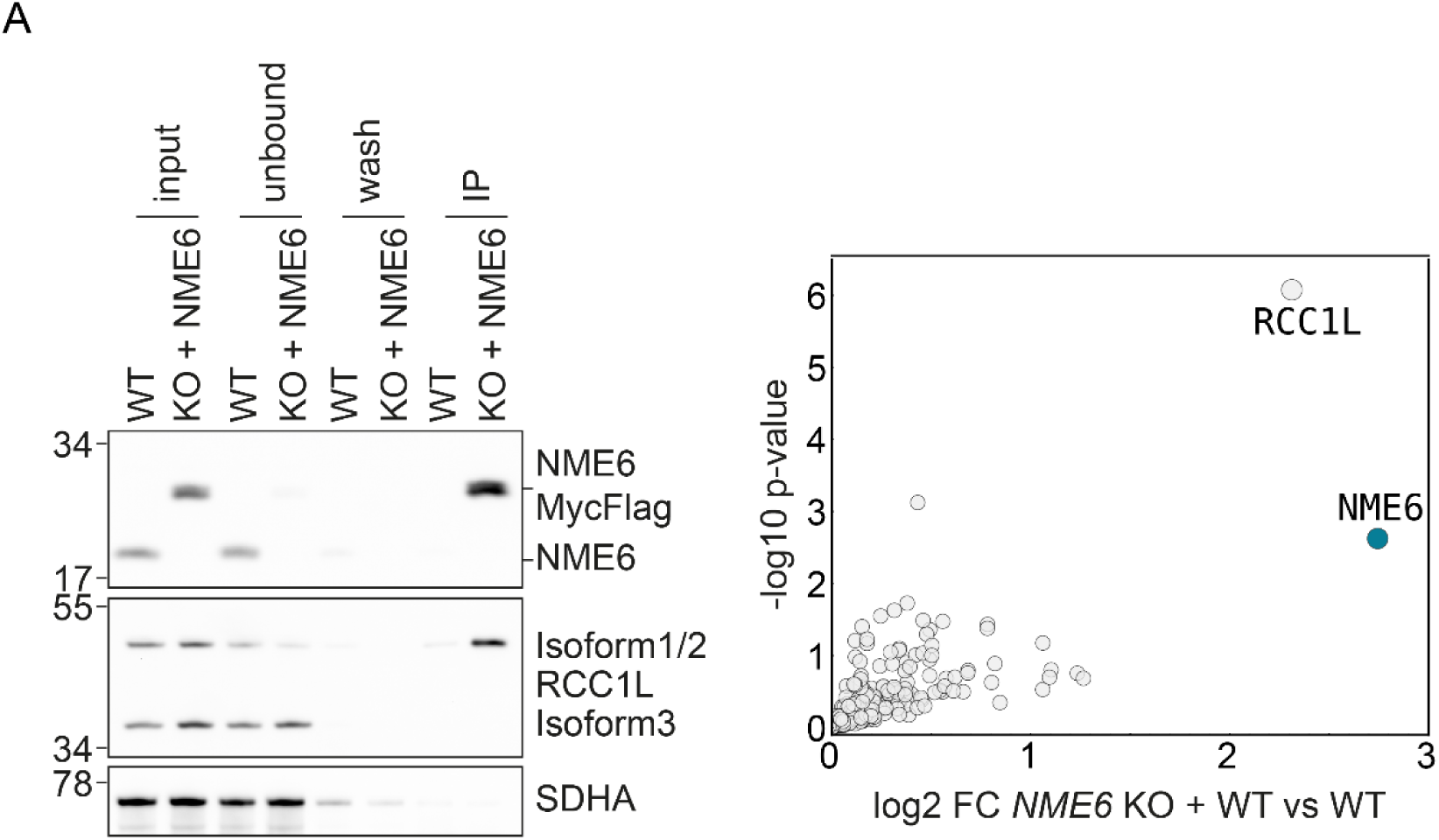
Proteomic analysis of immunoprecipitates shows interaction between NME6 and RCC1L. (A) Representative immunoblot (left) and proteomic analysis (right) following immunoprecipitation of NME6-MycFlag from HeLa cell mitochondrial lysates using a Flag antibody (n=4 independent experiments). FC, fold change.

**Figure S5.**
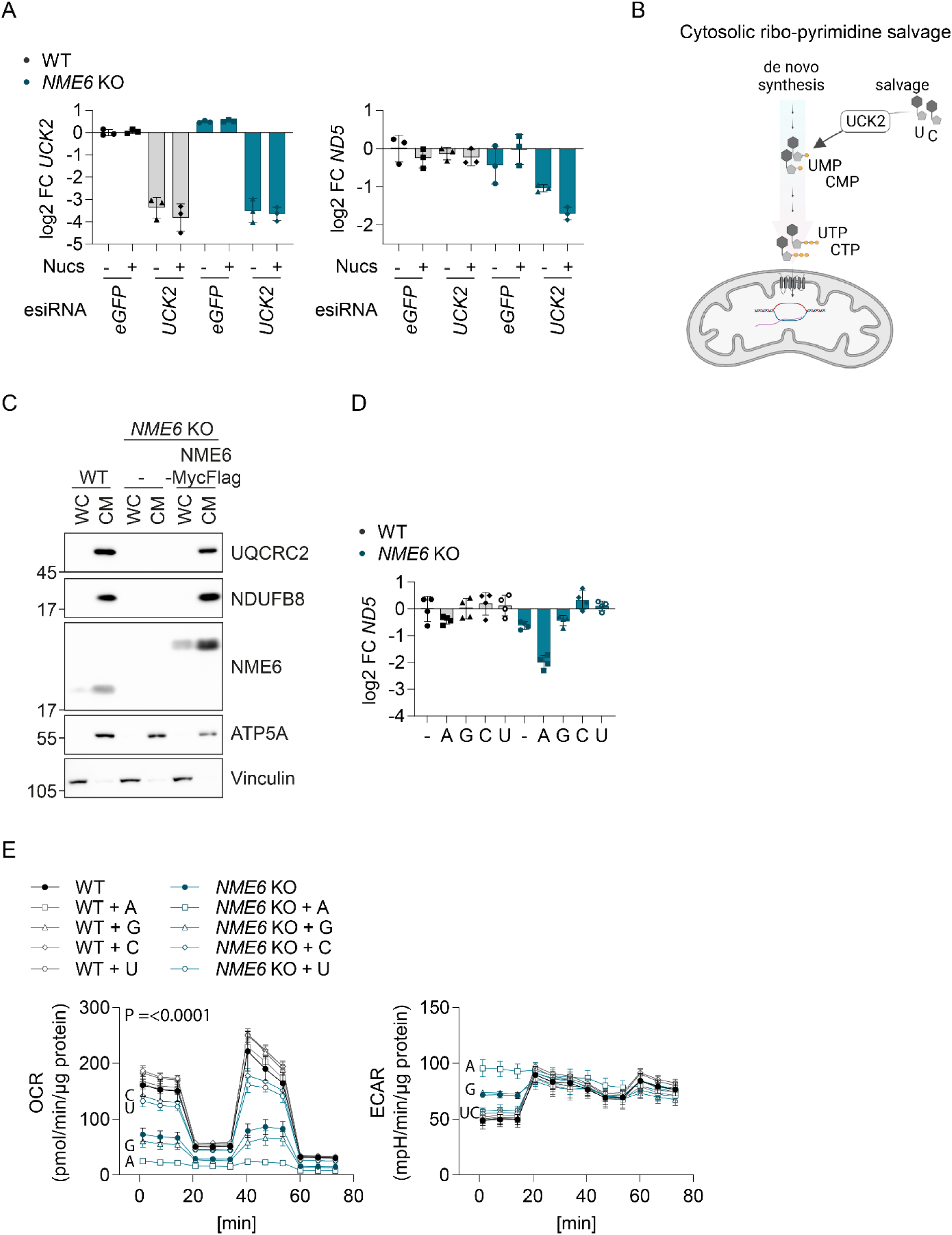
(related to Figure 5). NME6 supplies pyrimidines for mitochondrial transcription and OXPHOS. (A) *UCK2* and *ND5* transcript levels analysed by qRT-PCR in WT and *NME6* KO HeLa cells transfected with the indicated esiRNA and incubated with or without nucleosides (100 µM) for 72 h (log2; n=3 independent cultures from Fig. 3D). (B) Scheme to highlight the role played by UCK2 in pyrimidine nucleotide salvage. (C) Representative immunoblot showing the enrichment of mitochondrial proteins in crude mitochondrial fractions taken from the indicated HeLa cell lines during the isolation of mitochondria for LC-MS based metabolomics in Fig. 5F. WC, whole cell; CM, crude mitochondria. (D) *ND5* transcript levels analysed by qRT-PCR in WT and *NME6* KO HeLa cells incubated with the indicated nucleoside species for 48 h as in Fig. 5G (A, adenosine; G, guanosine; C, cytidine; U, uridine; log2 transformed; 100 µM; n=4 independent cultures). (E) Oxygen consumption rates (OCR) and extracellular acidification rates (ECAR) of WT and *NME6* KO HeLa cells incubated with or without individual nucleosides for a minimum of 120 h. Mitochondrial stress test was performed as in Fig. 3A (n=3 independent experiments). The basal and maximal respiration rates shown in Fig. 5H were determined from OCR measurements before injection of oligomycin and after injection of FCCP respectively (A, adenosine; G, guanosine; C, cytidine; U, uridine; 100 µM; n=3 independent cultures).

## Data availability

The mass spectrometry proteomics data have been deposited to the ProteomeXchange Consortium via the PRIDE [1] partner repository with the dataset identifier PXD038391^84^.

Statistical tests were performed with GraphPad Prism software version 9.4.1. on at least 3 independent cultures collected at different days.

## Materials and Methods

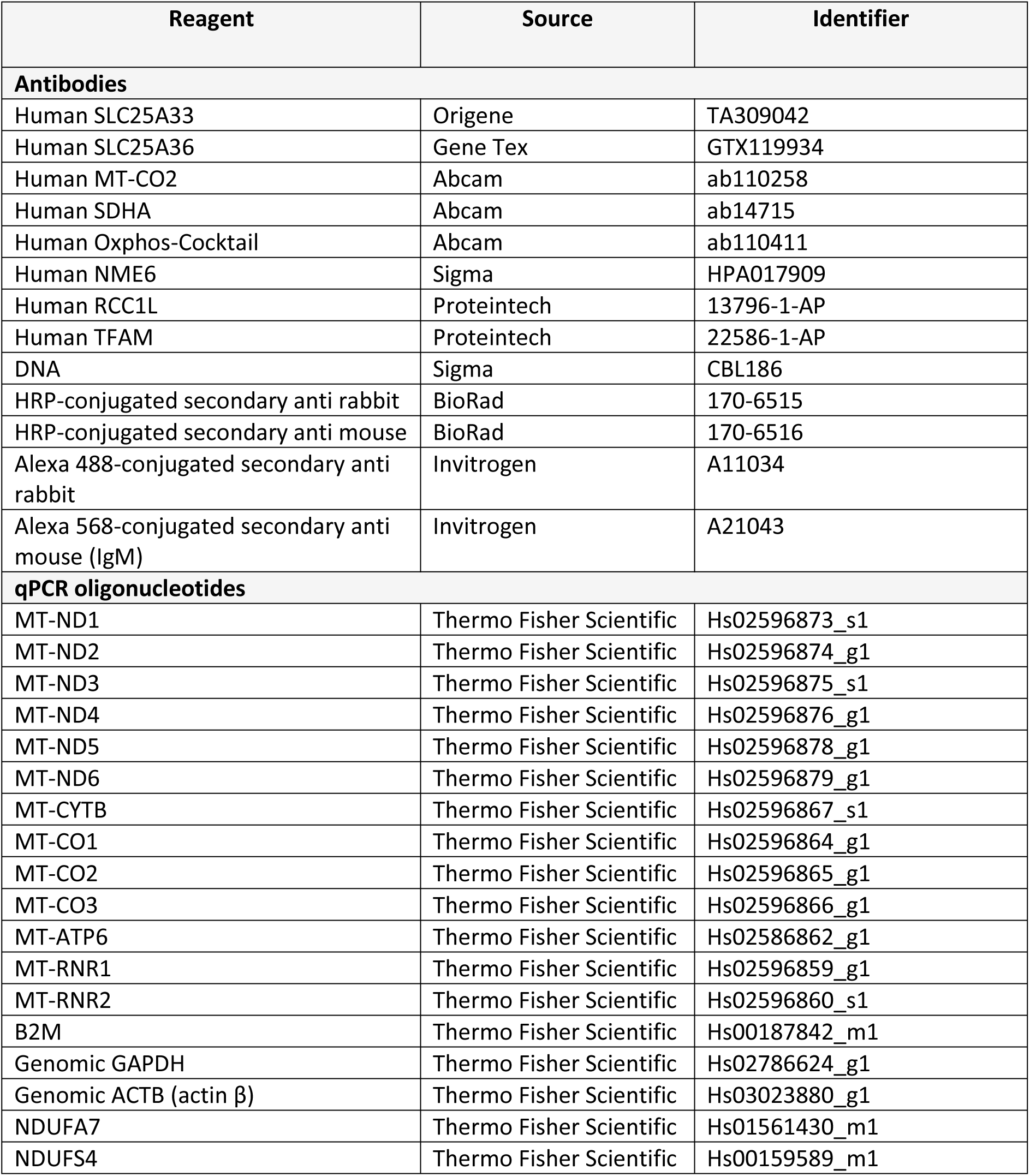
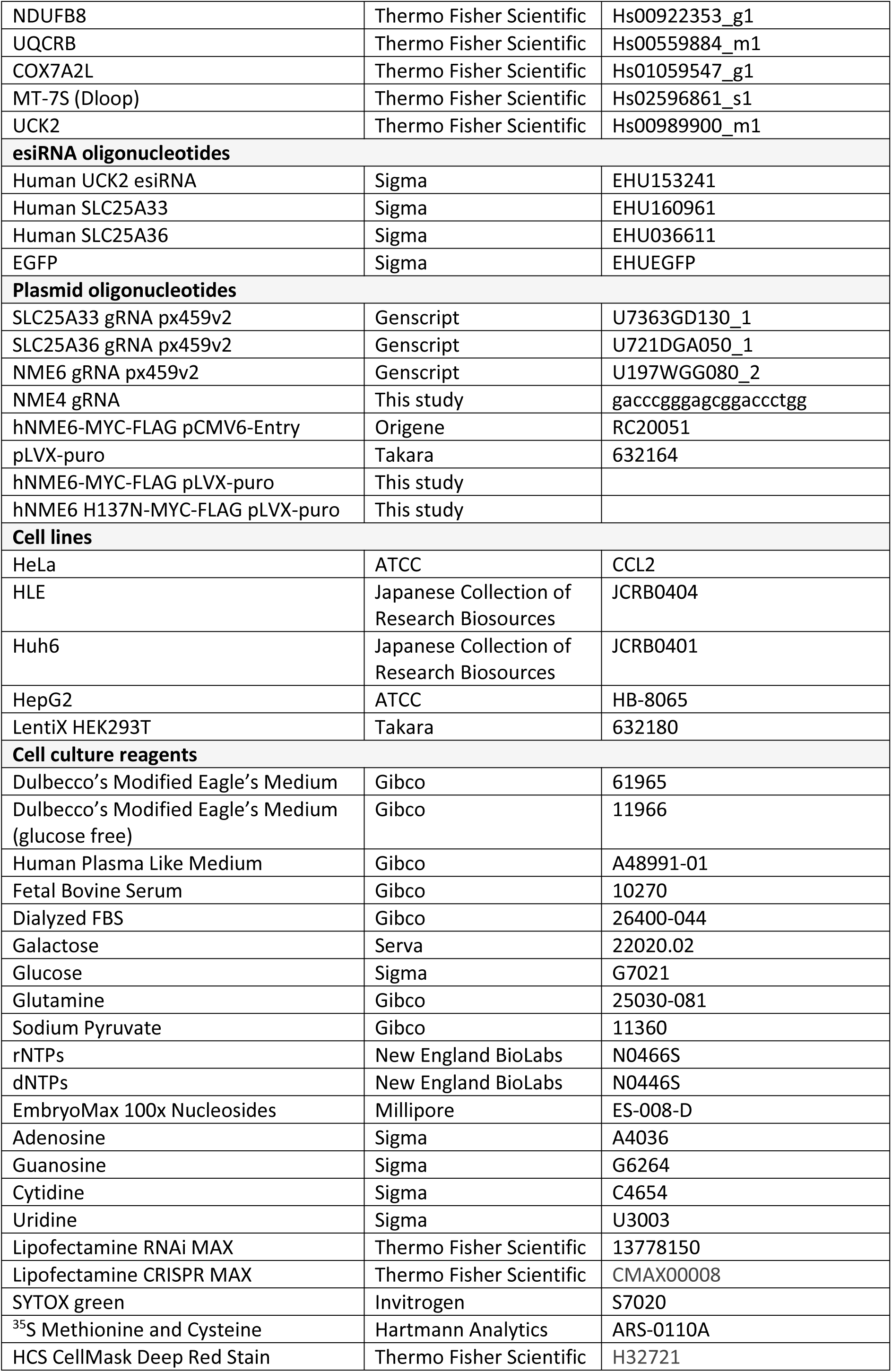
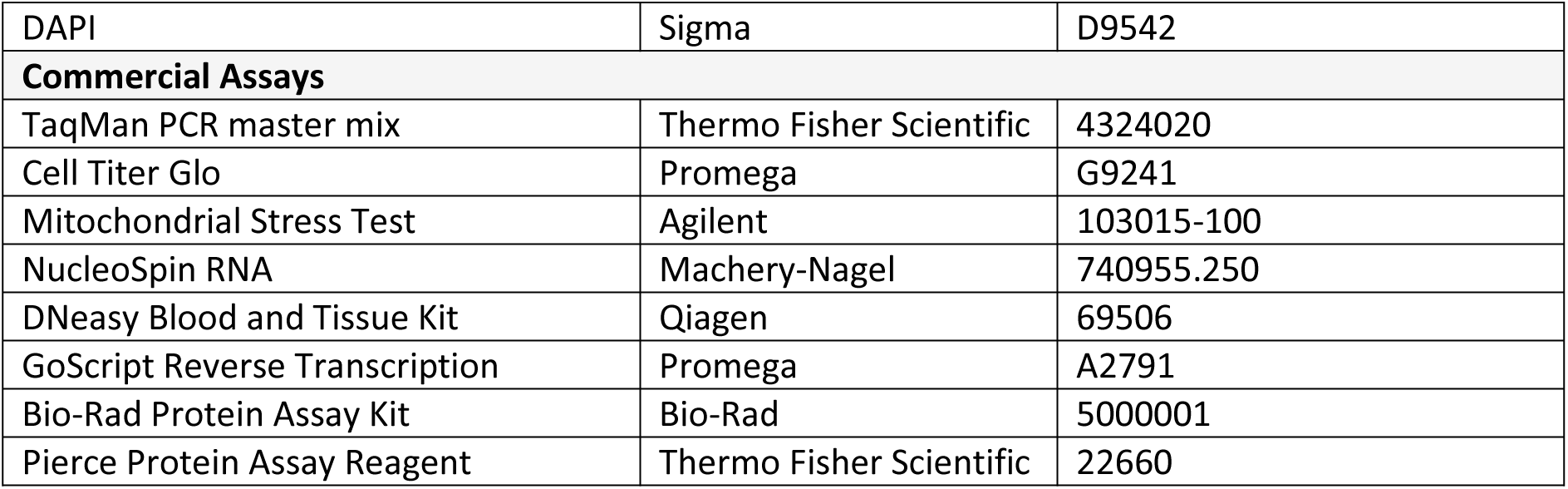

### Cell culture

HeLa, HLE, HepG2 and Huh6 cells were grown in Dulbecco’s Modified Eagle’s Medium (DMEM) containing 10 % fetal bovine serum (FBS) and maintained at 37 °C and 5 % CO_2_, if not stated otherwise. Alternatively, cells were cultured in either glucose free DMEM supplemented with 10 % FBS, 10 mM galactose and uridine (200 µg/ml), or in human plasma like medium (HPLM) supplemented with 10 % dialyzed FBS. All cultured cell lines were routinely tested for *Mycoplasma* contamination.

For supplementation experiments, nucleosides (100 µM), rNTPs (100 µM) or dNTPs (100 µM) were added to the medium for at least 48 h. RNA interference experiments were performed by reverse transfection of 2×10^5^ cells with 5 µg esiRNA using Lipofectamine RNAiMax.

### Generation of cell lines

Knockout (KO) cells were generated using CRISPR-SpCas9 mediated gene editing. SpCas9 and guide RNA (gRNA) were expressed using transient transfection of px459 v2 expression vector (Genscript). Polyclonal cultures were obtained by puromycin selection prior to monoclone selection by serial dilution. Polyclonal cultures and individual clones were validated by immunoblotting and genomic sequencing. HeLa and HLE NME6 KO cells expressing NME6-MycFlag or NME6 H137NMyc-Flag were generated by lentiviral transduction. Lenti-X HEK293T cells were transfected with either pLVX-NME6Myc-Flag or pLVX-NME6^H137N^ Myc-Flag using Lenti-X Packaging Single Shots (Takara). The viral supernatant was collected after 48 h, cleared from cell debris by centrifugation and added to HeLa NME6 KO cells together with polybrene (4 µg/ml). Virus-containing medium was removed after 24 h and puromycin selection (1 µg/ml) was started after an additional 24-48 h.

### Cell proliferation assays

HeLa cell proliferation was monitored by live cell imaging using the Incucyte S3 instrument (Sartorius). Image analysis was performed using Incucyte Software 2019 RevB. 5×10^3^ cells per well were seeded onto a 96 well plate and confluency was assessed every 6 h by phase contrast imaging until 100 % confluency was reached. Proliferation rates were determined from the slope of the exponential growth phase (24-72 h). Cell death was visualized by SYTOX green. SYTOX green was added to the assay medium (1:30000) and relative cell death was calculated by area of SYTOX green puncta divided by phase contrast cell area. Relative growth of HLE, Huh6 and HepG2 cells was determined on each day after 5×10^3^ cells were seeded per well of a 96-well plate. Total ATP luminescence was measured with the Cell Titer Glo viability assay (Promega) using a Glomax luminometer (Promega).

### DNA extraction, RNA extraction and cDNA synthesis

For mitochondrial DNA (mtDNA) measurements, genomic DNA was isolated for cell pellets using the Blood and Tissue DNA extraction kit (Qiagen). RNA was isolated from cell pellets using the RNA extraction kit (Macherey-Nagel) and 1-2 µg of RNA was reverse transcribed into cDNA using GoScript (Promega).

### Cell lysis and SDS-PAGE

Cells were collected in ice cold phosphate buffered saline (PBS). Cell pellets were lysed in RIPA buffer (50 mM Tris-HCl pH7.4, 150 mM NaCl, 1 % Triton X100, 0.5 % DOC, 0.1 % SDS and 1 mM EDTA) for 30 min at 4 °C. Lysates were cleared by centrifugation at 20000 xg for 10 min at 4 °C. Protein concentration was determined by Bradford assay (Bio-Rad). Protein lysates were mixed with 4x Laemmli buffer and analysed by 10 % SDS-PAGE and immunoblot.

### Quantitative PCR

For mtDNA measurements, 10 ng of genomic DNA were amplified using TaqMan PCR master mix (Thermo Fisher Scientific). MtDNA levels were assessed by the delta delta ct method using *MT-7S*, *CYTB* and *MT-ND6* as mitochondrial probes and *GAPDH* and *ACTB* as nuclear DNA controls. For the measurements of nuclear and mitochondrial transcripts,10 ng of cDNA were amplified using TaqMan PCR master mix. Expression levels were calculated by the delta delta ct method, for which *B2M* was used as control.

### Design of arrayed single guide RNA (sgRNA) library

The custom Mito Transporter and Salvage Pathway sgRNA library was purchased from Synthego and consisted of three sgRNA sequences designed to generate deletions in early exons of each target gene. The 116 target genes included all genes encoding proteins designated as “Small Molecule Transporters” in MitoCarta 3.0 ^47^ as well as putative mitochondrial metabolite carriers identified in a recent proteomic evaluation of mitochondria ^85^ and solute carriers with proposed mitochondrial localisation^86^. Mitochondrial pyrimidine salvage pathway genes were selected in addition to *TFAM* and controls, including non-targeting sgRNA (*NTC1*) and Polo-like kinase 1 (*PLK1*). All target genes and sgRNA sequences are listed in Table S1.

### Arrayed CRISPR-SpCas9 screen

The sgRNA library was reconstituted to 5 µM in Tris-EDTA (pH 8.0) and distributed to 96-well daughter plates. Note that the volumes in the following procedure correspond to individual transfections per well and were scaled up for the entire library in triplicate. Cas9 solution (0.5 µL SpCas9 2NLS nuclease, 1 µL Lipofectamine Cas9 plus reagent, 10.5 µL OptiMEM) was added to each well containing 1 µL sgRNA using the XRD-384 automated reagent dispenser (Fluidx). The Cas9-sgRNA mix was next stamped onto Cell Carrier Ultra 96-well plates (PerkinElmer) in triplicate prior to addition of the transfection reagent (0.35 µL Lipofectamine CRISPRMax, 10.15 uL OptiMEM) using the XRD-384. Each plate was placed on an orbital shaker at 300 rpm for 10 min at room temperature. In parallel, a cell suspension of SLC25A33/SLC25A36 DKO HeLa cells (clone #2) was prepared in DMEM + 10 % FBS (4×10^4^ cells/mL) and added (100 µL/ well) to the sgRNA:Cas9:LipofectamineCRISPRMax transfection mixture. Cells were distributed and incubated at 37 °C and 5 % CO_2_. After 6 h, the media and transfection mix was replaced with fresh media containing 150 µM uridine using a plate washer (BioTekELx405) to aspirate and XRD-384 to dispense. At 96 h post-transfection, the media was replaced with 80 µL of 4 % formaldehyde in DMEM for 10 mins. The cells were then washed twice in 150 µL PBS and permeabilised with 0.1 % Triton-TX100 in 80 µL PBS for 20 min prior to two further PBS washes. Primary antibody staining was performed sequentially with anti-TFAM (ProteinTech; 1:800) and anti-DNA (Sigma; 1:800) antibodies in 40 µL PBS for 30 min at room temperature. Secondary antibody staining was also performed sequentially with anti-rabbit IgG-Alexa 488 nm (Thermo Fisher Scientific; 1:1000) and anti-mouse IgM-Alexa 568 nm (Thermo Fisher Scientific; 1:1000) antibodies in 40 µL PBS for 30 min at room temperature. Each antibody staining was followed by three PBS washes. Finally, DAPI (Merk; 0.5µg/ml) and HCS CellMask Deep Red (Thermo Fisher Scientific; 1:20000) were combined in 200 µL PBS per well for 30mins prior to three final PBS washes. Plates were stored in the dark at 4°C with 200 µL PBS in each well prior to imaging.

Plates were imaged with an OperaPhenix High Content Analysis System (PerkinElmer) using a 20X objective (25 fields per well; single plane) and 63X water objective (24 fields per well; 5 Z-planes with 1 µm separation). Imaging was performed using 405 nm (DAPI), 488 nm (TFAM), 561 nm (DNA) and 640 nm (CellMask) excitation. Maximum intensity projections were generated in each channel using all planes and analysis was performed using Harmony 4.9 High-Content Imaging and Analysis Software (PerkinElmer). Nuclei masks were defined as DAPI positive structures above 30 um^2^ in area and were counted to determine cell number. The 20X objective was used to calculate cell number per well and all other analysis was performed with the 63X objective. MtDNA was measured within HCS CellMask stained cytoplasmic regions upon exclusion of the nuclei. MtDNA intensity was analysed using standard mean intensity and mtDNA puncta were identified and measured using the “Find Spots” algorithm (relative spot intensity: >0.045, splitting sensitivity: 1).

Across the three replicates, cell number was depleted by over 75 % in cells transfected with the lethal control sgRNA (PLK1) compared to non-targeting control sgRNA, which confirmed efficient transfection with Cas9:sgRNA ribonucleoprotein complexes in our screen (Table S1)

### Oxygen consumption and extracellular acidification measurements

Mitochondrial ATP-linked respiration and extra cellular acidification rate was measured by the Seahorse XFe96 Analyzer using the Mito Stress Test kit (Agilent). 4×10^4^ cells were seeded and grown for 24 h n DMEM containing 10 % FBS. For supplementation experiments, cells were cultured for at least 48 h in nucleoside containing medium prior to the experiment. Growth medium was exchanged to assay medium containing, glutamine, pyruvate and glucose. oligomycin (2 µM), FCCP (0.5 µM) and rotenone and antimycin A (0.5 µM each) injections were used to calculate basal respiration, ATP-linked ATP production and maximal respiration, respectively. Results were normalized to the amount of protein per well.

### Mitochondrial isolation

Cells were collected in ice cold PBS. Cell pellets were resuspended in 1 ml ice cold mitochondrial isolation buffer (containing 220 mM mannitol, 70 mM sucrose, 5 mM HEPES-KOH pH 7.4 and 1 mM EGTA-KOH + complete protease inhibitor). Cells were lysed detergent-free, using 10 strokes of a 1 ml syringe equipped with a 27g needle. Cell lysates were spun down at 600 xg for 5 min at 4 °C and the supernatant was separated from the remaining cell pellet. The supernatant was spun down for 10 min at 8000 xg at 4 °C. Mitochondria-enriched pellets were used for IP-, proteomic- and metabolomics experiments. Purity of the mitochondria-enriched fractions was verified by SDS-PAGE and immunoblot.

### Immunoprecipitation

Mitochondria-enriched pellets of HeLa WT and NME6 KO + NME6-MycFlag expressing cells (500 µg) were resuspended in 500 µl IP buffer (60 mM Tris-HCl and 300 mM KAc-KOH pH 7.4) Mitochondria were solubilized with digitonin (5 g/g protein) for 30 min at 4 °C while shaking on a ThermoMixer (shaking: 550 rpm). Mitochondrial lysates were spun down at 20000 xg for 15 min at 4 °C. The supernatant was mixed with Flag-agarose beads (Sigma) and incubated for 2 h at 4 °C. After 2 h, the supernatant was removed by centrifugation at 500 xg for 30 s and the remaining beads were washed three times with wash buffer (IP buffer containing 0.1 % digitonin). Bound proteins were eluted from the beads using 60 µl of 1x Laemmli buffer, samples were incubated for 10 min at 40 °C. The eluate was separated from the beads by centrifugation at 1000 xg for 3 min. Eluates were used for SDS-PAGE, immunoblot analysis and LC-MS based proteomics.

### Mitochondrial translation assay

Mitochondrial translation rates were monitored by incorporation rates of radioactive methionine and cysteine (^35^S) (Hartmann Analytic) into mitochondrial encoded proteins. Therefore, 3×10^5^ cells were cultured for 24 h. Growth medium was washed out and replaced with minimal medium depleted of methionine and cysteine. Cytosolic translation was blocked by emetine (100 µg/ml) for 30 min. ^35^S methionine and cysteine (50 µCi each) was added to the medium for 15-60 min. Cells were lysed and protein extracts were subjected to SDS-PAGE. Incorporation rates were visualized by autoradiography using a Typhoon FLA9500 imager (GE healthcare). Mitochondrial proteins were labelled according to their respective molecular weight.

### Extraction of polar metabolites

Cell pellets and mitochondrial pellets were resuspended in -20 °C cold extraction buffer (HPLC-grade ultrapure 40 % MeOH, 40 % acetonitrile, 20 % water and 0.1 µg/ml ^13^C labeled ATP). Pellets were dissolved by sonication at 4 °C followed by an incubation for 30 min at 4 °C while shaking at 1500 rpm. The metabolite containing supernatant was cleared by centrifugation at 20000 xg for 10 min and subsequently transferred to a speedvac concentrator to fully evaporate the extraction buffer. The remaining protein pellet from the extraction was used to determine protein concentration of the sample.

### Anion-Exchange Chromatography Mass Spectrometry (AEX-MS) for the analysis of nucleotides and deoxynucleotides

Extracted metabolites from crude- and mito-preparations were resuspended in 100 µl of UPLC/MS grade water (Biosolve) and transferred to p*olypropylene* autosampler vials (Chromatography Accessories Trott, Germany).

The samples were analysed using a Dionex ionchromatography system (Integrion Thermo Fisher Scientific) as described previously ^87^. In brief, 5 µL of polar metabolite extract were injected in push partial mode, using an overfill factor of 1, onto a Dionex IonPac AS11-HC column (2 mm × 250 mm, 4 μm particle size, Thermo Fisher Scientific) equipped with a Dionex IonPac AG11-HC guard column (2 mm × 50 mm, 4 μm, Thermo Fisher Scientific). The column temperature was held at 30 °C, while the auto sampler was set to 6 °C. A potassium hydroxide gradient was generated using a potassium hydroxide cartridge (Eluent Generator, Thermo Scientific), which was supplied with deionized water (Millipore). The metabolite separation was carried at a flow rate of 380 µL/min, applying the following gradient conditions: 0-3 min, 10 mM KOH; 3-12 min, 10−50 mM KOH; 12-19 min, 50-100 mM KOH; 19-22 min, 100 mM KOH, 22-23 min, 100-10 mM KOH. The column was re-equilibrated at 10 mM for 3 min.

For the analysis of metabolic pool sizes the eluting compounds were detected in negative ion mode using full scan measurements in the mass range m/z 77 – 770 on a Q-Exactive HF high resolution MS (Thermo Fisher Scientific). The heated electrospray ionization (HESI) source settings of the mass spectrometer were: Spray voltage 3.2 kV, capillary temperature was set to 300 °C, sheath gas flow 50 AU, aux gas flow 20 AU at a temperature of 330 °C and a sweep gas glow of 2 AU. The S-lens was set to a value of 60.

The semi-targeted LC-MS data analysis was performed using the TraceFinder software (Version 5.1, Thermo Fisher Scientific). The identity of each compound was validated by authentic reference compounds, which were measured at the beginning and the end of the sequence. For data analysis the area of the deprotonated [M-H^+^]^-1^ or doubly deprotonated [M-2H]^-2^ mono-isotopologue mass peaks of every required compound were extracted and integrated using a mass accuracy <3 ppm and a retention time tolerance of <0.05 min as compared to the independently measured reference compounds. These areas were then normalized to the internal standard, which was added to the extraction buffer, followed by a normalization to the protein content of the analyzed sample. Values were log2 transformed and normalized to the WT mean.

### Sample preparation for mass spectrometry-based proteomics

For whole proteome analysis, 60 µl of 4 % SDS in 100 mM HEPES-KOH (pH=8.5) was pre-heated to 70 °C and added to the cell pellet for further 10 min incubation at 70 °C on a ThermoMixer (shaking: 550 rpm). The protein concentration was determined using the 660 nm Protein Assay (Thermo Fisher Scientific, #22660). 20 µg of protein was subjected to tryptic digestion. For immunoprecipitation analysis, the LDS buffer eluate was directly used. Proteins were reduced (10 mM TCEP) and alkylated (20 mM CAA) in the dark for 45 min at 45 °C. Samples were subjected to an SP3-based digestion ^88^. Washed SP3 beads (Sera-Mag (TM) Magnetic Carboxylate Modified Particles (Hydrophobic, GE44152105050250), Sera-Mag(TM) Magnetic Carboxylate Modified Particles (Hydrophilic, GE24152105050250) from Sigma Aldrich) were mixed equally, and 3 µl of bead slurry were added to each sample. Acetonitrile was added to a final concentration of 50 % and washed twice using 70 % ethanol (V=200 µl) on an in-house made magnet. After an additional acetonitrile wash (V=200µl), 5 µl digestion solution (10 mM HEPES-KOH pH=8.5 containing trypsin (0.5 µg, Sigma) and LysC (0.5 µg, Wako)) was added to each sample and incubated overnight at 37 °C. Peptides were desalted on a magnet using 2 x 200 µl acetonitrile. Peptides were eluted in 10 µl 5 % DMSO in LC-MS water (Sigma Aldrich) in an ultrasonic bath for 10 min. Formic acid and acetonitrile were added to a final concentration of 2.5 % and 2 %, respectively. Samples were stored at -20 °C before subjection to LC-MS/MS analysis.

### Liquid chromatography and mass spectrometry

LC-MS/MS instrumentation consisted of an Easy-LC 1200 (Thermo Fisher Scientific) coupled via a nano-electrospray ionization source to an Exploris 480 mass spectrometer (Thermo Fisher Scientific, Bremen, Germany). An in-house packed column (inner diameter: 75 µm, length: 40 cm) was used for peptide separation. A binary buffer system (A: 0.1 % formic acid and B: 0.1 % formic acid in 80 % acetonitrile) was applied as follows:

Whole proteome analysis: Linear increase of buffer B from 4 % to 27 % within 70 min, followed by a linear increase to 45 % within 5 min. The buffer B content was further ramped to 65 % within 5 min and then to 95 % within 5 min. 95 % buffer B was kept for a further 5 min to wash the column.

Immunoprecipitation analysis: Linear increase of buffer B from 4 % to 27 % within 40 min, followed by a linear increase to 45 % within 5 min. The buffer B content was further ramped to 65 % within 5 min and then to 95 % within 5 min. 95 % buffer B was kept for a further 5 min to wash the column.

Prior to each sample, the column was washed using 5 µl buffer A and the sample was loaded using 8 µl buffer A.

The RF Lens amplitude was set to 55 %, the capillary temperature was 275 °C and the polarity was set to positive. MS1 profile spectra were acquired using a resolution of 120000 (at 200 m/z) at a mass range of 320-1150 m/z and an AGC target of 1 × 10^6^.

For MS/MS independent spectra acquisition, 48 equally spaced windows were acquired at an isolation m/z range of 15 Th, and the isolation windows overlapped by 1 Th. The fixed first mass was 200 m/z. The isolation center range covered a mass range of 357–1060 m/z. Fragmentation spectra were acquired at a resolution of 15000 at 200 m/z using a maximal injection time of 22 ms and stepped normalized collision energies (NCE) of 26, 28, and 30. The default charge state was set to 3. The AGC target was set to 3e6 (900 % - Exploris 480). MS2 spectra were acquired in centroid mode.

### Proteomics data analysis

DIA-NN (Data-Independent Acquisition by Neural Networks) v 1.8 ^89^ was used to analyse data-independent raw files. The spectral library was created using the reviewed-only Uniport reference protein (*Homo sapiens*, 20350 entries, downloaded September 2019) with the ‘Deep learning-based spectra and RTs prediction’ turned on. Protease was set to trypsin and a maximum of 1 miss cleavage was allowed. N-term M excision was set as a variable modification and carbamidomethylation at cysteine residues was set as a fixed modification. The peptide length was set to 7–30 amino acids and the precursor m/z range was defined from 340 – 1200 m/z. The option ‘Quantitative matrices’ was enabled. The FDR was set to 1 % and the mass accuracy (MS2 and MS1) as well as the scan window was set to 0 (automatic inference via DIA-NN). Match between runs (MBR) was enabled. The Neuronal network classifier worked in ‘double pass mode’ and protein interference was set to ‘Isoform IDs’. The quantification strategy was set to ‘robust LC (high accuracy)’ and cross-run normalization was defined as ‘RT-dependent’.

The ‘pg’ (protein group) output (MaxLFQ intensities ^90^) was further processed using Instant Clue ^91^ including and pairwise comparison using an unpaired two-sided t-test or one-way ANOVA followed by a permutation-based FDR correction (5 %).

MitoCarta 3.0 ^47^ and Uniprot-based Gene Ontology annotations were used for filtering. Hierarchical clustering, heatmaps and volcano plots were generated using the InstantClue software ^91^ v. 0.10.10.

## References

1. Winter, J.M., Yadav, T., and Rutter, J. (2022). Stressed to death: Mitochondrial stress responses connect respiration and apoptosis in cancer. Molecular Cell. 10.1016/j.molcel.2022.07.012.

2. Fernandez-Vizarra, E., and Zeviani, M. (2021). Mitochondrial disorders of the OXPHOS system. FEBS Letters 595, 1062–1106. https://doi.org/10.1002/1873-3468.13995.

3. Filograna, R., Mennuni, M., Alsina, D., and Larsson, N.G. (2021). Mitochondrial DNA copy number in human disease: the more the better? FEBS Lett 595, 976–1002. 10.1002/1873-3468.14021.

4. Russell, O.M., Gorman, G.S., Lightowlers, R.N., and Turnbull, D.M. (2020). Mitochondrial Diseases: Hope for the Future. Cell 181, 168–188. https://doi.org/10.1016/j.cell.2020.02.051.

5. Vafai, S.B., and Mootha, V.K. (2012). Mitochondrial disorders as windows into an ancient organelle. Nature 491, 374–383. 10.1038/nature11707.

6. Gustafsson, C.M., Falkenberg, M., and Larsson, N.-G. (2016). Maintenance and Expression of Mammalian Mitochondrial DNA. Annual Review of Biochemistry 85, 133–160. 10.1146/annurev-biochem-060815-014402.

7. D’Souza, A.R., and Minczuk, M. (2018). Mitochondrial transcription and translation: overview. Essays Biochem 62, 309–320. 10.1042/ebc20170102.

8. Lane, A.N., and Fan, T.W. (2015). Regulation of mammalian nucleotide metabolism and biosynthesis. Nucleic Acids Res 43, 2466–2485. 10.1093/nar/gkv047.

9. Wang, L. (2016). Mitochondrial purine and pyrimidine metabolism and beyond. Nucleosides Nucleotides Nucleic Acids 35, 578–594. 10.1080/15257770.2015.1125001.

10. Mathews, C.K., and Song, S. (2007). Maintaining precursor pools for mitochondrial DNA replication. The FASEB Journal 21, 2294–2303. https://doi.org/10.1096/fj.06-7977rev.

11. Ferraro, P., Pontarin, G., Crocco, L., Fabris, S., Reichard, P., and Bianchi, V. (2005). Mitochondrial deoxynucleotide pools in quiescent fibroblasts: a possible model for mitochondrial neurogastrointestinal encephalomyopathy (MNGIE). J Biol Chem 280, 24472–24480. 10.1074/jbc.M502869200.

12. Suomalainen, A., and Isohanni, P. (2010). Mitochondrial DNA depletion syndromes – Many genes, common mechanisms. Neuromuscular Disorders 20, 429–437. https://doi.org/10.1016/j.nmd.2010.03.017.

13. Saada, A., Shaag, A., Mandel, H., Nevo, Y., Eriksson, S., and Elpeleg, O. (2001). Mutant mitochondrial thymidine kinase in mitochondrial DNA depletion myopathy. Nat Genet 29, 342–344. 10.1038/ng751.

14. Hamalainen, R.H., Landoni, J.C., Ahlqvist, K.J., Goffart, S., Ryytty, S., Rahman, M.O., Brilhante, V., Icay, K., Hautaniemi, S., Wang, L.Y., et al. (2019). Defects in mtDNA replication challenge nuclear genome stability through nucleotide depletion and provide a unifying mechanism for mouse progerias. Nature Metabolism 1, 958-+. 10.1038/s42255-019-0120-1.

15. Sprenger, H.-G., MacVicar, T., Bahat, A., Fiedler, K.U., Hermans, S., Ehrentraut, D., Ried, K., Milenkovic, D., Bonekamp, N., Larsson, N.-G., et al. (2021). Cellular pyrimidine imbalance triggers mitochondrial DNA–dependent innate immunity. Nature Metabolism 3, 636–650. 10.1038/s42255-021-00385-9.

16. Zhong, Z., Liang, S., Sanchez-Lopez, E., He, F., Shalapour, S., Lin, X.J., Wong, J., Ding, S., Seki, E., Schnabl, B., et al. (2018). New mitochondrial DNA synthesis enables NLRP3 inflammasome activation. Nature 560, 198–203. 10.1038/s41586-018-0372-z.

17. Ernst, O., Sun, J., Lin, B., Banoth, B., Dorrington, M.G., Liang, J., Schwarz, B., Stromberg, K.A., Katz, S., Vayttaden, S.J., et al. (2021). A genome-wide screen uncovers multiple roles for mitochondrial nucleoside diphosphate kinase D in inflammasome activation. Sci Signal 14. 10.1126/scisignal.abe0387.

18. Zhu, X., Xie, X., Das, H., Tan, B.G., Shi, Y., Al-Behadili, A., Peter, B., Motori, E., Valenzuela, S., Posse, V., et al. (2022). Non-coding 7S RNA inhibits transcription via mitochondrial RNA polymerase dimerization. Cell. https://doi.org/10.1016/j.cell.2022.05.006.

19. Zhou, T., Sang, Y.H., Cai, S., Xu, C., and Shi, M.H. (2021). The requirement of mitochondrial RNA polymerase for non-small cell lung cancer cell growth. Cell Death Dis 12, 751. 10.1038/s41419-021-04039-2.

20. Wan, L., Wang, Y., Zhang, Z., Wang, J., Niu, M., Wu, Y., Yang, Y., Dang, Y., Hui, S., Ni, M., et al. (2021). Elevated TEFM expression promotes growth and metastasis through activation of ROS/ERK signaling in hepatocellular carcinoma. Cell Death Dis 12, 325. 10.1038/s41419-021-03618-7.

21. Fei, Z.Y., Wang, W.S., Li, S.F., Zi, J.J., Yang, L., Liu, T., Ao, S., Liu, Q.Q., Cui, Q.H., Yu, M., and Xiong, W. (2020). High expression of the TEFM gene predicts poor prognosis in hepatocellular carcinoma. J Gastrointest Oncol 11, 1291–1304. 10.21037/jgo-20-120.

22. Bonekamp, N.A., Peter, B., Hillen, H.S., Felser, A., Bergbrede, T., Choidas, A., Horn, M., Unger, A., Di Lucrezia, R., Atanassov, I., et al. (2020). Small-molecule inhibitors of human mitochondrial DNA transcription. Nature 588, 712–716. 10.1038/s41586-020-03048-z.

23. Favre, C., Zhdanov, A., Leahy, M., Papkovsky, D., and O’Connor, R. (2010). Mitochondrial pyrimidine nucleotide carrier (PNC1) regulates mitochondrial biogenesis and the invasive phenotype of cancer cells. Oncogene 29, 3964–3976. 10.1038/onc.2010.146.

24. Di Noia, M.A., Todisco, S., Cirigliano, A., Rinaldi, T., Agrimi, G., Iacobazzi, V., and Palmieri, F. (2014). The Human SLC25A33 and SLC25A36 Genes of Solute Carrier Family 25 Encode Two Mitochondrial Pyrimidine Nucleotide Transporters*. Journal of Biological Chemistry 289, 33137–33148. https://doi.org/10.1074/jbc.M114.610808.

25. Floyd, S., Favre, C., Lasorsa, F.M., Leahy, M., Trigiante, G., Stroebel, P., Marx, A., Loughran, G., O’Callaghan, K., Marobbio, C.M., et al. (2007). The insulin-like growth factor-I-mTOR signaling pathway induces the mitochondrial pyrimidine nucleotide carrier to promote cell growth. Mol Biol Cell 18, 3545–3555. 10.1091/mbc.e06-12-1109.

26. Shahroor, M.A., Lasorsa, F.M., Porcelli, V., Dweikat, I., Di Noia, M.A., Gur, M., Agostino, G., Shaag, A., Rinaldi, T., Gasparre, G., et al. (2022). PNC2 (SLC25A36) Deficiency Associated With the Hyperinsulinism/Hyperammonemia Syndrome. J Clin Endocrinol Metab 107, 1346–1356. 10.1210/clinem/dgab932.

27. Van Dyck, E., Jank, B., Ragnini, A., Schweyen, R.J., Duyckaerts, C., Sluse, F., and Foury, F. (1995). Overexpression of a novel member of the mitochondrial carrier family rescues defects in both DNA and RNA metabolism in yeast mitochondria. Mol Gen Genet 246, 426–436. 10.1007/bf00290446.

28. Parisi, M.A., and Clayton, D.A. (1991). Similarity of human mitochondrial transcription factor 1 to high mobility group proteins. Science 252, 965–969. 10.1126/science.2035027.

29. Garrido, N., Griparic, L., Jokitalo, E., Wartiovaara, J., van der Bliek, A.M., and Spelbrink, J.N. (2003). Composition and dynamics of human mitochondrial nucleoids. Mol Biol Cell 14, 1583–1596. 10.1091/mbc.e02-07-0399.

30. Legros, F.d.r., Malka, F., Frachon, P., Lombès, A., and Rojo, M. (2004). Organization and dynamics of human mitochondrial DNA. Journal of Cell Science 117, 2653–2662. 10.1242/jcs.01134.

31. Ekstrand, M.I., Falkenberg, M., Rantanen, A., Park, C.B., Gaspari, M., Hultenby, K., Rustin, P., Gustafsson, C.M., and Larsson, N.-G. (2004). Mitochondrial transcription factor A regulates mtDNA copy number in mammals. Human Molecular Genetics 13, 935–944. 10.1093/hmg/ddh109.

32. Larsson, N.-G., Wang, J., Wilhelmsson, H., Oldfors, A., Rustin, P., Lewandoski, M., Barsh, G.S., and Clayton, D.A. (1998). Mitochondrial transcription factor A is necessary for mtDNA maintance and embryogenesis in mice. Nature Genetics 18, 231–236. 10.1038/ng0398-231.

33. Kanki, T., Ohgaki, K., Gaspari, M., Gustafsson, C.M., Fukuoh, A., Sasaki, N., Hamasaki, N., and Kang, D. (2004). Architectural Role of Mitochondrial Transcription Factor A in Maintenance of Human Mitochondrial DNA. Molecular and Cellular Biology 24, 9823–9834. doi:10.1128/MCB.24.22.9823-9834.2004.

34. Bonekamp, N.A., Jiang, M., Motori, E., Garcia Villegas, R., Koolmeister, C., Atanassov, I., Mesaros, A., Park, C.B., and Larsson, N.-G. (2021). High levels of TFAM repress mammalian mitochondrial DNA transcription in vivo. Life Science Alliance 4, e202101034. 10.26508/lsa.202101034.

35. Feric, M., Sarfallah, A., Dar, F., Temiakov, D., Pappu, R.V., and Misteli, T. (2022). Mesoscale structure–function relationships in mitochondrial transcriptional condensates. Proceedings of the National Academy of Sciences 119, e2207303119. doi:10.1073/pnas.2207303119.

36. West, A.P., Khoury-Hanold, W., Staron, M., Tal, M.C., Pineda, C.M., Lang, S.M., Bestwick, M., Duguay, B.A., Raimundo, N., MacDuff, D.A., et al. (2015). Mitochondrial DNA stress primes the antiviral innate immune response. Nature 520, 553–557. 10.1038/nature14156.

37. Boissan, M., Schlattner, U., and Lacombe, M.L. (2018). The NDPK/NME superfamily: state of the art. Lab Invest 98, 164–174. 10.1038/labinvest.2017.137.

38. Chen, C.-W., Wang, H.-L., Huang, C.-W., Huang, C.-Y., Lim, W.K., Tu, I.-C., Koorapati, A., Hsieh, S.-T., Kan, H.-W., Tzeng, S.-R., et al. (2019). Two separate functions of NME3 critical for cell survival underlie a neurodegenerative disorder. Proceedings of the National Academy of Sciences 116, 566–574. doi:10.1073/pnas.1818629116.

39. Tokarska-Schlattner, M., Boissan, M., Munier, A., Borot, C., Mailleau, C., Speer, O., Schlattner, U., and Lacombe, M.-L. (2008). The Nucleoside Diphosphate Kinase D (NM23-H4) Binds the Inner Mitochondrial Membrane with High Affinity to Cardiolipin and Couples Nucleotide Transfer with Respiration*. Journal of Biological Chemistry 283, 26198–26207. https://doi.org/10.1074/jbc.M803132200.

40. Milon, L., Meyer, P., Chiadmi, M., Munier, A., Johansson, M., Karlsson, A., Lascu, I., Capeau, J., Janin, J., and Lacombe, M.-L. (2000). The Human nm23-H4 Gene Product Is a Mitochondrial Nucleoside Diphosphate Kinase*. Journal of Biological Chemistry 275, 14264–14272. https://doi.org/10.1074/jbc.275.19.14264.

41. Proust, B., Radic, M., Vidacek, N.S., Cottet, C., Attia, S., Lamarche, F., Ackar, L., Mikulcic, V.G., Tokarska-Schlattner, M., Cetkovic, H., et al. (2021). NME6 is a phosphotransfer-inactive, monomeric NME/NDPK family member and functions in complexes at the interface of mitochondrial inner membrane and matrix. Cell Biosci 11, 195. 10.1186/s13578-021-00707-0.

42. Tsuiki, H., Nitta, M., Furuya, A., Hanai, N., Fujiwara, T., Inagaki, M., Kochi, M., Ushio, Y., Saya, H., and Nakamura, H. (1999). A novel human nucleoside diphosphate (NDP) kinase, Nm23-H6, localizes in mitochondria and affects cytokinesis. J Cell Biochem 76, 254–269. 10.1002/(sici)1097-4644(20000201)76:2<254::aid-jcb9>3.0.co;2-g.

43. Schlattner, U. (2021). The Complex Functions of the NME Family-A Matter of Location and Molecular Activity. Int J Mol Sci 22. 10.3390/ijms222313083.

44. Reitzer, L.J., Wice, B.M., and Kennell, D. (1979). Evidence that glutamine, not sugar, is the major energy source for cultured HeLa cells. Journal of Biological Chemistry 254, 2669–2676. https://doi.org/10.1016/S0021-9258(17)30124-2.

45. Rossignol, R., Gilkerson, R., Aggeler, R., Yamagata, K., Remington, S.J., and Capaldi, R.A. (2004). Energy Substrate Modulates Mitochondrial Structure and Oxidative Capacity in Cancer Cells. Cancer Research 64, 985–993. 10.1158/0008-5472.Can-03-1101.

46. Rossiter, N.J., Huggler, K.S., Adelmann, C.H., Keys, H.R., Soens, R.W., Sabatini, D.M., and Cantor, J.R. (2021). CRISPR screens in physiologic medium reveal conditionally essential genes in human cells. Cell Metab. 10.1016/j.cmet.2021.02.005.

47. Rath, S., Sharma, R., Gupta, R., Ast, T., Chan, C., Durham, T.J., Goodman, R.P., Grabarek, Z., Haas, M.E., Hung, W.H.W., et al. (2021). MitoCarta3.0: an updated mitochondrial proteome now with sub-organelle localization and pathway annotations. Nucleic Acids Res 49, D1541–D1547. 10.1093/nar/gkaa1011.

48. Ahola, S., Rivera Mejías, P., Hermans, S., Chandragiri, S., Giavalisco, P., Nolte, H., and Langer, T. (2022). OMA1-mediated integrated stress response protects against ferroptosis in 50 mitochondrial cardiomyopathy. Cell Metab 34, 1875–1891.e1877. 10.1016/j.cmet.2022.08.017.

49. Mick, E., Titov, D.V., Skinner, O.S., Sharma, R., Jourdain, A.A., and Mootha, V.K. (2020). Distinct mitochondrial defects trigger the integrated stress response depending on the metabolic state of the cell. Elife 9. 10.7554/eLife.49178.

50. Wang, T., Yu, H., Hughes, N.W., Liu, B., Kendirli, A., Klein, K., Chen, W.W., Lander, E.S., and Sabatini, D.M. (2017). Gene Essentiality Profiling Reveals Gene Networks and Synthetic Lethal Interactions with Oncogenic Ras. Cell 168, 890–903.e815. 10.1016/j.cell.2017.01.013.

51. Amici, D.R., Jackson, J.M., Truica, M.I., Smith, R.S., Abdulkadir, S.A., and Mendillo, M.L. (2021). FIREWORKS: a bottom-up approach to integrative coessentiality network analysis. Life Sci Alliance 4. 10.26508/lsa.202000882.

52. Floyd, Brendan J., Wilkerson, Emily M., Veling, Mike T., Minogue, Catie E., Xia, C., Beebe, Emily T., Wrobel, Russell L., Cho, H., Kremer, Laura S., Alston, Charlotte L., et al. (2016). Mitochondrial Protein Interaction Mapping Identifies Regulators of Respiratory Chain Function. Molecular Cell 63, 621–632. https://doi.org/10.1016/j.molcel.2016.06.033.

53. Antonicka, H., Lin, Z.Y., Janer, A., Aaltonen, M.J., Weraarpachai, W., Gingras, A.C., and Shoubridge, E.A. (2020). A High-Density Human Mitochondrial Proximity Interaction Network. Cell Metab 32, 479–497 e479. 10.1016/j.cmet.2020.07.017.

54. Reyes, A., Favia, P., Vidoni, S., Petruzzella, V., and Zeviani, M. (2020). RCC1L (WBSCR16) isoforms coordinate mitochondrial ribosome assembly through their interaction with GTPases. PLoS Genet 16, e1008923. 10.1371/journal.pgen.1008923.

55. Miranda, M., Bonekamp, N.A., and Kuhl, I. (2022). Starting the engine of the powerhouse: mitochondrial transcription and beyond. Biol Chem. 10.1515/hsz-2021-0416.

56. Tan, B.G., Mutti, C.D., Shi, Y., Xie, X., Zhu, X., Silva-Pinheiro, P., Menger, K.E., Díaz-Maldonado, H., Wei, W., Nicholls, T.J., et al. (2022). The human mitochondrial genome contains a second light strand promoter. Mol Cell. 10.1016/j.molcel.2022.08.011.

57. Lascu, I., and Gonin, P. (2000). The Catalytic Mechanism of Nucleoside Diphosphate Kinases. Journal of Bioenergetics and Biomembranes 32, 237–246. 10.1023/A:1005532912212.

58. Zimmermann, H. (1999). Two novel families of ectonucleotidases: molecular structures, catalytic properties and a search for function. Trends in Pharmacological Sciences 20, 231–236. 10.1016/S0165-6147(99)01293-6.

59. Pastor-Anglada, M., and Perez-Torras, S. (2018). Emerging Roles of Nucleoside Transporters. Front Pharmacol 9, 606. 10.3389/fphar.2018.00606.

60. Pooler, A.M., Guez, D.H., Benedictus, R., and Wurtman, R.J. (2005). Uridine enhances neurite outgrowth in nerve growth factor-differentiated PC12 [corrected]. Neuroscience 134, 207–214. 10.1016/j.neuroscience.2005.03.050.

61. Szczepanowska, K., and Trifunovic, A. (2021). Tune instead of destroy: How proteolysis keeps OXPHOS in shape. Biochimica et Biophysica Acta (BBA) - Bioenergetics 1862, 148365. https://doi.org/10.1016/j.bbabio.2020.148365.

62. Deshwal, S., Fiedler, K.U., and Langer, T. (2020). Mitochondrial Proteases: Multifaceted Regulators of Mitochondrial Plasticity. Annual Review of Biochemistry 89, 501–528. 10.1146/annurev-biochem-062917-012739.

63. Dogan, S.A., Pujol, C., Maiti, P., Kukat, A., Wang, S., Hermans, S., Senft, K., Wibom, R., Rugarli, E.I., and Trifunovic, A. (2014). Tissue-specific loss of DARS2 activates stress responses independently of respiratory chain deficiency in the heart. Cell Metab 19, 458–469. 10.1016/j.cmet.2014.02.004.

64. Buchet, K., and Godinot, C. (1998). Functional F1-ATPase Essential in Maintaining Growth and Membrane Potential of Human Mitochondrial DNA-depleted ρ° Cells*. Journal of Biological Chemistry 273, 22983–22989. https://doi.org/10.1074/jbc.273.36.22983.

65. Edenberg, Ellen R., Downey, M., and Toczyski, D. (2014). Polymerase Stalling during Replication, Transcription and Translation. Current Biology 24, R445–R452. https://doi.org/10.1016/j.cub.2014.03.060.

66. Ruzzenente, B., Metodiev, M.D., Wredenberg, A., Bratic, A., Park, C.B., Cámara, Y., Milenkovic, D., Zickermann, V., Wibom, R., Hultenby, K., et al. (2012). LRPPRC is necessary for polyadenylation and coordination of translation of mitochondrial mRNAs. The EMBO Journal 31, 443–456. https://doi.org/10.1038/emboj.2011.392.

67. Gohil, V.M., Nilsson, R., Belcher-Timme, C.A., Luo, B., Root, D.E., and Mootha, V.K. (2010). Mitochondrial and Nuclear Genomic Responses to Loss of LRPPRC Expression*. Journal of Biological Chemistry 285, 13742–13747. https://doi.org/10.1074/jbc.M109.098400.

68. Wagner, A., Hofmeister, O., Rolland, S.G., Maiser, A., Aasumets, K., Schmitt, S., Schorpp, K., Feuchtinger, A., Hadian, K., Schneider, S., et al. (2019). Mitochondrial Alkbh1 localizes to mtRNA granules and its knockdown induces the mitochondrial UPR in humans and C. elegans. J Cell Sci 132. 10.1242/jcs.223891.

69. Zaganelli, S., Rebelo-Guiomar, P., Maundrell, K., Rozanska, A., Pierredon, S., Powell, C.A., Jourdain, A.A., Hulo, N., Lightowlers, R.N., Chrzanowska-Lightowlers, Z.M., et al. (2017). The Pseudouridine Synthase RPUSD4 Is an Essential Component of Mitochondrial RNA Granules. J Biol Chem 292, 4519–4532. 10.1074/jbc.M116.771105.

70. Antonicka, H., and Shoubridge, Eric A. (2015). Mitochondrial RNA Granules Are Centers for Posttranscriptional RNA Processing and Ribosome Biogenesis. Cell Reports 10, 920–932. https://doi.org/10.1016/j.celrep.2015.01.030.

71. Antonicka, H., Choquet, K., Lin, Z.Y., Gingras, A.C., Kleinman, C.L., and Shoubridge, E.A. (2017). A pseudouridine synthase module is essential for mitochondrial protein synthesis and cell viability. EMBO Rep 18, 28–38. 10.15252/embr.201643391.

72. Xavier, V.J., and Martinou, J.-C. (2021). RNA Granules in the Mitochondria and Their Organization under Mitochondrial Stresses. Int J Mol Sci 22, 9502.

73. Kühl, I., Miranda, M., Posse, V., Milenkovic, D., Mourier, A., Siira, S.J., Bonekamp, N.A., Neumann, U., Filipovska, A., Polosa, P.L., et al. (2016). POLRMT regulates the switch between replication primer formation and gene expression of mammalian mtDNA. Sci Adv 2, e1600963. 10.1126/sciadv.1600963.

74. Falkenberg, M. (2018). Mitochondrial DNA replication in mammalian cells: overview of the pathway. Essays Biochem 62, 287–296. 10.1042/ebc20170100.

75. Gandhi, V.V., and Samuels, D.C. (2011). Enzyme kinetics of the mitochondrial deoxyribonucleoside salvage pathway are not sufficient to support rapid mtDNA replication. PLoS Comput Biol 7, e1002078. 10.1371/journal.pcbi.1002078.

76. Besse, A., Wu, P., Bruni, F., Donti, T., Graham, B.H., Craigen, W.J., McFarland, R., Moretti, P., Lalani, S., Scott, K.L., et al. (2015). The GABA transaminase, ABAT, is essential for mitochondrial nucleoside metabolism. Cell Metab 21, 417–427. 10.1016/j.cmet.2015.02.008.

77. Zhong, F., Liang, S., and Zhong, Z. (2019). Emerging Role of Mitochondrial DNA as a Major Driver of Inflammation and Disease Progression. Trends in Immunology 40, 1120–1133. https://doi.org/10.1016/j.it.2019.10.008.

78. Xu, Y., Johansson, M., and Karlsson, A. (2008). Human UMP-CMP Kinase 2, a Novel Nucleoside Monophosphate Kinase Localized in Mitochondria*. Journal of Biological Chemistry 283, 1563–1571. https://doi.org/10.1074/jbc.M707997200.

79. Boissan, M., Montagnac, G., Shen, Q., Griparic, L., Guitton, J., Romao, M., Sauvonnet, N., Lagache, T., Lascu, I., Raposo, G., et al. (2014). Membrane trafficking. Nucleoside diphosphate kinases fuel dynamin superfamily proteins with GTP for membrane remodeling. Science 344, 1510–1515. 10.1126/science.1253768.

80. Lacombe, M.L., Tokarska-Schlattner, M., Boissan, M., and Schlattner, U. (2018). The mitochondrial nucleoside diphosphate kinase (NDPK-D/NME4), a moonlighting protein for cell homeostasis. Lab Invest 98, 582–588. 10.1038/s41374-017-0004-5.

81. Pontarin, G., Ferraro, P., Håkansson, P., Thelander, L., Reichard, P., and Bianchi, V. (2007). p53R2-dependent ribonucleotide reduction provides deoxyribonucleotides in quiescent human fibroblasts in the absence of induced DNA damage. J Biol Chem 282, 16820–16828. 10.1074/jbc.M701310200.

82. Nusinow, D.P., Szpyt, J., Ghandi, M., Rose, C.M., McDonald, E.R., III, Kalocsay, M., Jané-Valbuena, J., Gelfand, E., Schweppe, D.K., Jedrychowski, M., et al. (2020). Quantitative Proteomics of the Cancer Cell Line Encyclopedia. Cell 180, 387–402.e316. 10.1016/j.cell.2019.12.023.

83. Smith, J.C., and Sheltzer, J.M. (2022). Genome-wide identification and analysis of prognostic features in human cancers. Cell Rep 38, 110569. 10.1016/j.celrep.2022.110569.

84. Perez-Riverol, Y., Bai, J., Bandla, C., García-Seisdedos, D., Hewapathirana, S., Kamatchinathan, S., Kundu, Deepti J., Prakash, A., Frericks-Zipper, A., Eisenacher, M., et al. (2021). The PRIDE database resources in 2022: a hub for mass spectrometry-based proteomics evidences. Nucleic Acids Research 50, D543–D552. 10.1093/nar/gkab1038.

85. Morgenstern, M., Peikert, C.D., Lübbert, P., Suppanz, I., Klemm, C., Alka, O., Steiert, C., Naumenko, N., Schendzielorz, A., Melchionda, L., et al. (2021). Quantitative high-confidence human mitochondrial proteome and its dynamics in cellular context. Cell Metabolism 33, 2464–2483.e2418. https://doi.org/10.1016/j.cmet.2021.11.001.

86. Meixner, E., Goldmann, U., Sedlyarov, V., Scorzoni, S., Rebsamen, M., Girardi, E., and Superti-Furga, G. (2020). A substrate-based ontology for human solute carriers. Mol Syst Biol 16, e9652. 10.15252/msb.20209652.

87. Schwaiger, M., Rampler, E., Hermann, G., Miklos, W., Berger, W., and Koellensperger, G. (2017). Anion-Exchange Chromatography Coupled to High-Resolution Mass Spectrometry: A Powerful Tool for Merging Targeted and Non-targeted Metabolomics. Anal Chem 89, 7667–7674. 10.1021/acs.analchem.7b01624.

88. Hughes, C.S., Foehr, S., Garfield, D.A., Furlong, E.E., Steinmetz, L.M., and Krijgsveld, J. (2014). Ultrasensitive proteome analysis using paramagnetic bead technology. Molecular Systems Biology 10. 10.15252/msb.20145625.

89. Demichev, V., Messner, C.B., Vernardis, S.I., Lilley, K.S., and Ralser, M. (2020). DIA-NN: neural networks and interference correction enable deep proteome coverage in high throughput. Nature Methods 17, 41-+. 10.1038/s41592-019-0638-x.

90. Cox, J., Hein, M.Y., Luber, C.A., Paron, I., Nagaraj, N., and Mann, M. (2014). Accurate proteome-wide label-free quantification by delayed normalization and maximal peptide ratio extraction, termed MaxLFQ. Mol Cell Proteomics 13, 2513–2526. 10.1074/mcp.M113.031591.

91. Nolte, H., MacVicar, T.D., Tellkamp, F., and Kruger, M. (2018). Instant Clue: A Software Suite for Interactive Data Visualization and Analysis. Sci Rep 8, 12648. 10.1038/s41598-018-31154-6.

